# Using correlative and mechanistic species distribution models to predict vector-borne disease risk for the current and future environmental and climatic change: a case study of West Nile Virus in the UK

**DOI:** 10.1101/2024.09.12.612656

**Authors:** A. J. Withers, S. Croft, R. Budgey, D. Warren, N. Johnson

## Abstract

Globally, vector-borne diseases have significant impacts on both animal and human health, and these are predicted to increase with the effects of climate change. Understanding the drivers of such diseases can help inform surveillance and control measures to minimise risks both now and in the future. In this study, we illustrate a generalised approach for assessing disease risk combining species distribution models of vector and wildlife hosts with data on livestock and human populations using the potential emergence of West Nile Virus (WNV) in the UK as a case study. Currently absent in the UK, WNV is an orthoflavivirus with a natural transmission cycle between *Culex* mosquitos (*Cx. pipiens* and *Cx. modestus*) and birds. It can spread into non-target hosts (e.g., equids, humans) via mosquito bites where it can cause febrile disease with encephalitis and mortality in severe cases. We compared six correlative species distribution models and selected the most appropriate for each vector based on a selection of performance measures and compared this to mechanistic species distribution models and known distributions. We then combined these with correlative species distribution models of representative avian hosts, equines, and human population data to predict risk of WNV occurrence. Our findings highlighted areas at greater risk of WNV due to higher habitat suitability for both avian hosts and vectors, and considered how this risk could change by 2100 under a best-case Shared Socioeconomic Pathway (SSP1) and worst-case (SSP5) future climate scenario. Generally, WNV risk in the future was found to increase in south-eastern UK and decrease further north. Overall, this paper presents how current and future vector distributions can be modelled and combined with projected host distributions to predict areas at greater risk of novel diseases. This is important for policy decision making and contingency preparedness to enable adaptation to changing environments and the resulting shifts in vector-borne diseases that are predicted to occur.

## Introduction

Vector-borne diseases have a significant impact upon global animal and human health, accounting for around 17% of all infectious diseases in humans, leading to over 700,000 deaths a year worldwide (World Health Organisation, 2020). Additionally, impacts on the livestock industry can result in devastating economic losses, as well as contributing to food shortages due to limited productivity (Faburay, 2015). Most insect vector species, including mosquitos, ticks, and biting midges, are exotherms meaning that their lifecycle and geographic ranges are highly influenced by the climate and environment. Consequently, the risk from the diseases they carry is expected to be significantly affected by future climate change and urbanisation, with range shifts predicted to occur globally (Endo and Amarasekare, 2022, Rocklöv and Dubrow, 2020, Noll et al., 2023). Understanding these potential range shifts is important for surveillance so mitigation measures can be developed to reduce impact on both human and animal health. For the UK, the risk of vector-borne disease is predicted to increase with the potential for increased globalisation and warming temperatures leading to a higher number of invasive mosquito species introductions and favourable conditions that could support year-round vector survival, leading to outbreaks of diseases such as dengue, chikungunya, and West Nile fever (Medlock and Leach, 2015, Townroe and Callaghan, 2014).

West Nile Virus (WNV) is an orthoflavivirus with a natural transmission cycle between mosquitos and birds. On rare occasions, mosquitos will spread WNV into non-target, dead-end hosts, such as humans and equids. Whilst WNV is usually asymptomatic in these non-target hosts, it can become symptomatic (e.g., headache, fever, rash) in around 20% of cases, with 1 in 150 human cases becoming neuroinvasive (e.g., acute aseptic meningitis, encephalitis) which can occasionally lead to severe illness or death (Pradier et al., 2012, World Health Organisation, 2017). In recent years, outbreaks of WNV have occurred in Europe in areas where the disease has not been present historically (e.g., France, Germany), and predictions suggest WNV could reach the UK (Vogels et al., 2017, Constant et al., 2022, Bessell et al., 2016). The most likely route of WNV introduction to the UK is via migrating birds from endemic regions of mainland Europe (Bessell et al., 2016). Some birds migrate to the UK from areas in Africa where WNV is present, however, the time taken for this migration to be completed means that viremia will have declined and transmission will no longer be possible when the birds arrive at their final destination (Pradier et al., 2012).

As well as significant effects on human and animal health, WNV can have severe economic impacts. For example, predictions of the economic impact in Belgium were around €4,500 per human case that led to hospitalisation which could be thousands during an outbreak (Humblet et al., 2016). As well as additional costs to the equestrian sector of around €17 million to vaccinate horses in at-risk areas to prevent further economic damage (Humblet et al., 2016). Therefore, understanding the distribution of WNV vectors and hosts in the UK can reveal where the risk is higher, this is important for contingency planning and mitigation to reduce the impact of an outbreak by enabling measures to be put in place to protect human and animal health, and consequently reduce economic stress (Medlock and Leach, 2015, Medlock et al., 2018, Gandy et al., 2022, Lawson et al., 2022).

There are two primary vectors of WNV in the UK, these are *Culex pipiens* f. *pipiens* (hereafter *Cx. pipiens*) and *Culex modestus*. *Cx. pipiens* has a widespread distribution, and mostly feeds on birds but has been known to feed on mammals so could potentially spread WNV into non-target hosts (e.g., humans, equids). C*x. modestus* is an invasive species in the UK that was first discovered in the south of England (Portsmouth) in the 1940’s, then rediscovered in the north Kent marshes in southeast England in 2003 where it had established (Golding et al., 2012, Vaux et al., 2015). Since then, multiple other records have occurred across the UK, but the distribution of *Cx. modestus* is currently much more limited than that of *Cx. pipiens*. *Cx. modestus* is of interest as a key bridge vector for WNV transmission and host spillover as it is known to bite both mammals and birds, especially in areas where frequent overlap of these potential blood meals occurs (Rudolf et al., 2021, Hernández-Triana et al., 2020, Soto and Delang, 2023).

When modelling the potential range of insect distributions there are two main approaches; correlative and mechanistic (Koch, 2021). Correlative species distribution models have been widely applied to predict distributions across all taxa by relating occurrence data to a set of environmental variables to determine suitability and then using the suitability model to predict species distributions across the whole study area (Koch, 2021, Elith et al., 2010, Hijmans and Elith, 2013, Valavi et al., 2021). There are a broad range of correlative species distribution models available, such as Random Forest (RF), Boosted Regression Tree (BRT), Support Vector Machine (SVM), Neural Network (NET) and MaxEnt (MAX). Each of these approaches have previously been used to produce species distribution models for a large range of taxa, including mosquitos, and each can outperform the other depending on the species and presence or absence data available (European Centre for Disease Prevention and Control, 2019, Beeman et al., 2021, Simons et al., 2019, Furlong et al., 2022, Senay et al., 2013, Senay and Worner, 2019, Richman et al., 2018, Gorris et al., 2021). It is rarely clear which individual model is the most appropriate. Ensemble approaches have been used to remedy this issue and can be beneficial, allowing multiple outputs to be combined into a single consensus model. However, determining how to combine different models is difficult and many approaches have been suggested (Simons et al., 2019, European Centre for Disease Prevention and Control, 2019, Beeman et al., 2021).

The correlative approaches discussed above each require presence-absence data (i.e., known absences) or presence-background (i.e., characterise where a species is less likely to be found based on known presences). Presence-absence data is often unavailable for species distribution modelling as this requires a consistent sampling effort over a widespread area to ensure that absences are true absences and not due to sampling bias (Elith et al., 2006, VanDerWal et al., 2009, Lobo et al., 2010, Iturbide et al., 2015, Croft and Smith, 2019). Therefore, presence-background is the most common approach for species distribution modelling, however, this requires artificially generating pseudo-absence or background points to represent areas where the species is absent. It can be difficult to determine background data for species with limited presence records, especially if it is an invasive species as they may not have reached all suitable locations within the area yet (Elith et al., 2010, Giovanelli et al., 2010). Whilst this is true of many species it requires extra consideration for insects as host and range expansions can occur rapidly in some species due to short life-cycles and multiple generations each year, meaning that the predicted distribution of insects may be underestimated if background points are created in areas where survival is possible (Koch, 2021). Conversely, mechanistic models do not rely on sampling data as they are based on *a priori* knowledge of a species and its response to climatic variables, including factors such as seasonal population growth, environmental stress, and survival constraints (Koch, 2021, Sutherst and Maywald, 1985, Kriticos et al., 2015). This means that they are usually limited to plants and insects where dependencies are more climatic so laboratory studies to measure tolerances (e.g., temperature, drought) can inform the models (Sutherst and Maywald, 1985, Poutsma et al., 2008, Taylor and Kumar, 2012, Khormi and Kumar, 2014, Kriticos et al., 2015, Early et al., 2022). A common mechanistic modelling approach is CLIMEX, which predicts a species’ potential range based on biological constraints (Sutherland et al., 2021, Kriticos et al., 2015). Previously, CLIMEX modelling has successfully been used to determine potential distributions of invasive mosquitos including *Aedes aegypti* and *Aedes atropalpus* (Khormi and Kumar, 2014, Scholte et al., 2009), and it can be beneficial for species with limited presence-absence data as it does not rely on sampling data (Sutherst and Maywald, 1985, Taylor and Kumar, 2012, Khormi and Kumar, 2014, Kriticos et al., 2015, Early et al., 2022).

Here, an approach of producing disease risk maps based on species distribution modelling of vector and host habitat suitability is presented, using WNV emergence in the UK as a case study. Correlative species distribution models are used to predict the range of each vector (*Cx. pipiens* and *Cx. modestus*), and a mechanistic species distribution model was also produced for each vector to further validate the correlative approaches. These models were then used to predict distributions across the UK for the current climate as well as future predictions to the year 2100 for two shared socioeconomic pathways; SSP1 (climate protection measures and development compatible with target of 2°C) and SSP5 (worst case scenario without climate measures and mitigations) (Fick and Hijmans, 2017). These individual vector maps were then combined to create an overall vector risk map for all climate scenarios. Following this, correlative species distribution models were produced of representative avian hosts for corvid and non-corvid Passeriformes. Once all current and future suitability predictions had been produced for vectors and hosts, these were then combined with horse or human density data to produce overall WNV risk maps for the current climate (based on horse or human density) and future environment (based on human density).

## Methods

### Model extent

Both vector and avian host models were conducted at a global extent, with a spatial-resolution of 30 arc-seconds (0.00833°), which equates to about 1km^2^ at the equator. The models were then used to predict distribution across the whole UK at the same resolution 0.00833°.

### Presence and absence data

#### Vector presence

Global species presence data for all confirmed records of *Culex pipiens* and *Culex modestus* were downloaded from the National Biodiversity Network (NBN) Atlas and the Global Biodiversity Information Facility (GBIF) for all time up to the download date of 7^th^ November 2022 (NBN Atlas, 2022, GBIF Org User, 2022). An additional presence record for *Cx. modestus* was included from a recent paper reporting the first recording of the species in Finland (Culverwell and Vapalahti, 2023). For each species, all presences were combined and cleaned in R (v. 4.3.3) to remove duplicated records and invalid points (e.g., over the sea) (CoordinateCleaner::*clean_coordinates*, v3.0.1, Zizka et al. (2019)). Presence points were then thinned based on environmental conditions prior to modelling to further reduce sampling bias (flexsdm::*occ_*filt, v1.3.4, Velazco et al. (2022)) (Castellanos et al., 2019).

#### Host presence

A wide variety of avian hosts can be susceptible to and spread WNV, however, the Passerine order are generally considered the most susceptible (Ziegler et al., 2022, Montecino-Latorre and Barker, 2018, Durand et al., 2010, Bergmann et al., 2023, Komar et al., 2003, Centers for Disease Control and Prevention, 2021). Here, we considered common species susceptible to WNV frequently observed around areas that may be suitable for mosquito vectors and humans or horses to act as representative avian hosts (Brugman et al., 2017). Selected avian host species were aggregated into two groups, corvids consisting of the common raven (*Corvus corax*), rook (*Corvus frugilegus*) and carrion crow (*Corvus corone*) and non-corvids comprising of the house sparrow (*Passer domesticus*), blackbird (*Turdus merula*) and barn swallow (*Hirundo rustica*).

Bird presence records for our selected species were taken from GBIF for April to October from 2013 to 2023 on 4^th^ June 2024 for all six avian host species included in these models (GBIF. Org User, 2024, GBIF.Org User, 2024d, GBIF.Org User, 2024e, GBIF.Org User, 2024b, GBIF.Org User, 2024a, GBIF.Org User, 2024c). GBIF was used for presences as it includes eBird observations, alongside many other submitters, thereby maximising the number of presence records available. These presences were cleaned using clean_coordinates in R (v.4.4.0) to remove duplicated records and invalid points (e.g., over the sea) (CoordinateCleaner::*clean_coordinates*, v3.0.1, Zizka et al. (2019)). Presence points were thinned based on environmental conditions prior to modelling to further reduce sampling bias (flexsdm::*occ_*filt, v1.3.4, Velazco et al. (2022)) (Castellanos et al., 2019).

#### Predictor variable selection

Twenty-one environmental predictor variables were selected for vectors and hosts for the current and future environmental conditions based on two Shared Socioeconomic Pathways (SSP), SSP126 and SSP585 (Table 1). For vectors and hosts a global extent was used, with a resolution of 0.00833, 0.00833 (i.e., about 1km^2^). To reduce multicollinearity in models Pearson correlation coefficient was used to remove climatic variables with a >0.7 correlation coefficient irrespective of presence or absence points (i.e., correlation across all predictors) to minimise the effects of spatial autocorrelation (*vifcor,* usdm, (Naimi et al., 2014)), and a dendrogram of clusters of correlates is available in Supplementary Information (SI) 1. Following this, the remaining variables included were bio2 (mean diurnal range), bio8 (mean temperature of the wettest quarter), bio9 (mean temperature of the driest quarter), bio14 (precipitation of the driest month), bio15 (precipitation seasonality), bio18 (precipitation of the warmest quarter), bio19 (precipitation of the coldest quarter), land use and human population density.

**Table 1:** The predictor variables used in modelling vectors and hosts. Each predictor variable was downloaded for the current time and for two Shared-Socioeconomic Pathways (SSP) in the future (up to 2100), these were SSP1 and SSP5. Those variables in italics were maintained in the model following variable selection based on correlation.

| Variable | Resolution | Source |
| --- | --- | --- |
| Bioclimatic variables based on UK-ESM1-0-LL:<br>• BIO1 = Annual Mean Temperature<br>• *BIO2 = Mean Diurnal Range*<br>• BIO3 = Isothermality<br>• BIO4 = Temperature Seasonality<br>• BIO5 = Max Temperature of Warmest Month<br>• BIO6 = Min Temperature of Coldest Month<br>• BIO7 = Temperature Annual Range<br>• *BIO8 = Mean Temperature of Wettest Quarter*<br>• *BIO9 = Mean Temperature of Driest Quarter*<br>• BIO10 = Mean Temperature of Warmest Quarter<br>• BIO11 = Mean Temperature of Coldest Quarter<br>• BIO12 = Annual Precipitation<br>• BIO13 = Precipitation of Wettest Month<br>• *BIO14 = Precipitation of Driest Month*<br>• *BIO15 = Precipitation Seasonality*<br>• BIO16 = Precipitation of Wettest Quarter<br>• BIO17 = Precipitation of Driest Quarter<br>• *BIO18 = Precipitation of Warmest Quarter*<br>• *BIO19 = Precipitation of Coldest Quarter* | 0.00833,<br>0.00833<br>(approximately<br>1km<sup>2</sup>) | (Fick and Hijmans, 2017) |
| *Land use (7 land types):* Forest, Grassland, Barren, Cropland, Urban, Water, Permanent snow and ice | 1km<sup>2</sup> (re-projected for modelling to 0.00833, 0.00833) | (Chen et al., 2022) |
| *Human population density* | 1km<sup>2</sup> (re-projected for modelling to 0.00833, 0.00833) | (Gao, 2020, Gao, 2017) |

#### Modelling approaches

For vectors, seven different modelling approaches were used, six of these were correlative approaches and one was a mechanistic approach used to determine suitability based on the biological constraints of *Cx. pipiens* and *Cx. modestus*. For hosts, four correlative approaches were used as CLIMEX is most appropriate for species strongly limited by climate due to factors such as temperature and moisture, whereas avian distribution is influenced strongly by additional factors. Mechanistic modelling was carried out using CLIMEX (version 4, Kriticos et al. (2015)). All correlative species distribution modelling and the production of figures was carried out in R (v 4.4.0).

### Correlative species distribution models for vectors and hosts

#### Vectors

Based on the presence data available following filtering, a model calibration area was determined based on a 100km buffer around presence points for *Cx. pipiens* and *Cx. modestus*, this radius was selected following trials of models using different buffers (5km, 50km, and 100km) and comparing performance across several different performance measures (full details are provided in Supplementary Information (SI) 2). Within this calibration area we randomly allocate 4716 and 144 background points (dismo::*randomPoints,* v.1.3-14, Hijmans et al. (2023)) for *Cx. pipiens* and *Cx. modestus* respectively to act as pseudo-absences. This method has the fewest assumptions about the presence data. The number of background points was selected as three-times the number of presences available as large numbers of absences have been shown to not be beneficial for species with small presence datasets, with fewer absences leading to increased accuracy in machine learning models (Liu et al., 2019, Barbet-Massin et al., 2012, Thuiller et al., 2024).

Prior to model fitting, the presence and background points datasets were partitioned into two datasets, one for training and one for testing. For vectors, this was based on the spatial autocorrelation of points based on Moran’s I, environmental similarity, and the number of points available to ensure a balanced division of points between the training and testing datasets (flexsdm::*part_sband,* v1.3.4, Velazco et al. (2022)) (Roberts et al., 2017, Santini et al., 2021, Velazco et al., 2022). Using a structured approach to partition a large database into training and testing datasets can help to overcome the sampling and spatial bias that can later affect performance measures and the evaluation of models in partitioning is carried out randomly (Valavi et al., 2022, Elith et al., 2006, Valavi et al., 2021, Roberts et al., 2017, Elith et al., 2010). Following data partitioning based on spatial autocorrelation, 1319 presence and 3064 background points were in the training dataset and 253 presence and 1531 background points were in the testing datasets for *Culex pipiens*. For *Cx. modestus,* 27 presence and 72 background points were in the training dataset and 21 presence and 70 background points were in the testing datasets.

The correlative modelling approaches used were Random Forest (RF, flexsdm::*tune_raf, v1.3.4, Velazco et al. (2022)*), Maxent (MAX, flexsdm::*tune_max, v1.3.4, Velazco et al. (2022)*), Generalized Boosted Regression (GBM, flexsdm::*tune_gbm, v1.3.4, Velazco et al. (2022)*), Neural Network (NET, flexsdm::*tune_net, v1.3.4, Velazco et al. (2022)*), Support Vector Machine (SVM, flexsdm::*tune_svm, v1.3.4, Velazco et al. (2022)*) and an Ensemble model based on all five individual models (flexsdm::fit_ensemble*, v1.3.4, Velazco et al. (2022))*.

All individual correlative models were fit using the *tune* functions in the *flexsdm* package, with selections based on three different thresholds, which were when sensitivity and specificity were equal (*equal_sens_spec*), when the sum of sensitivity and specificity was highest (*max_sens_spec*) and when the Sorensen threshold was highest (*max_sorensen*). A number of performance metrics were calculated during model fitting and considered for model evaluation, as well as visual comparison to known presence locations (European Centre for Disease Prevention and Control and European Food Safety Authority, 2022, GBIF Org User, 2022). A number of measures were used as there is often variation in how each model is scored, and determining suitability on a single measure can lead to poor model selection due to inherent, undetectable biases, such as model overfitting and sampling bias in the data used to build the models (Velazco et al., 2022, Valavi et al., 2022, Valavi et al., 2021, Hijmans and Elith, 2013, Roberts et al., 2017). The performance metrics used were True positive rate (TPR), True negative rate (TNR), Sorensen, Jaccard, F-measure of presence-background (FPB), Omission rate (OR), True skill statistic (TSS), Area under curve (AUC), Boyce index and Inverse mean absolute error (IMAE) (Velazco et al., 2022, Valavi et al., 2022, Valavi et al., 2021, Hijmans and Elith, 2013, Roberts et al., 2017). The TPR and TNR are the proportion of observed presences (TPR) or absences (TNR) in the test data that are correctly predicted by the model, and are sometimes referred to as sensitivity and specificity respectively. Several performance metrics are based on the TPR and TNR alongside the falsely classified presences (False Positive Rate; FPR) and absences (False Negative Rate; FNR) that give an indication of model performance, including the Sorensen and Jaccard similarity indices, FPB, OR and TSS. The Sorensen and Jaccard similarity indices are scored on a scale of 0 to 1, where 0 is a very poor similarity and 1 high similarity and thus indicates a better performing model. They are calculated very similarly, however, Sorensen assigns more importance to true positives by weighting them higher than true negatives in the index. The FPB is on a scale of 0 to 2, where a higher score is indicative of better model performance based on combining the TPR, TNR, FPR and FNR. The OR is the proportion of presences correctly assigned by the model. The TSS is calculated by combining the TPR and TNR to give an overall indication of performance, and is also on a scale of 0 to 1 where the closer to 1 the score is, the better the model is. The AUC is widely used in species distribution modelling to compare the TPR and FPR to assess model performance on a scale of 0 to 1, where 0.5 suggests the model is no better than random and 0.75 is generally considered good. The Boyce index measures how predictions made using the test dataset differ from a random distribution of the observed presences, and is scored on a scale of -1 to 1, where 1 indicates a good model and 0 suggests the model is no different to random. Negative values suggest that the predictions are frequently incorrect (i.e., presences located in areas predicted to be unsuitable). The IMAE considers the total number of correctly assigned presences and absences in relation to the total number of observations, and it is also given on a scale of 0 to 1, where 1 reflects a model performing well with no errors. Full details on how these performance measures are calculated are available in the *flexsdm* R package (Velazco et al., 2022).

#### Hosts

Using the presence data available following filtering for each individual species, a model calibration area was determined based on a convex polygon around presence points to represent the accessible area as this has previously been shown to improve model performance and worked well for other bird species distribution models (Rojas-Soto et al., 2024, Palacio et al., 2021, Rousseau and Betts, 2022). Background points were randomly allocated within the defined model calibration area (dismo::*randomPoints,* v.1.3-14, Hijmans et al. (2023)) for each species to act as pseudo-absences as this method has the fewest assumptions about the presence data. Due to the large number of presence data available, an equal number of background points was created (Liu et al., 2019, Barbet-Massin et al., 2012, Thuiller et al., 2024). Any datapoints outside the extent of environmental predictors were excluded from the dataset. Prior to model fitting, the species presence and background data were partitioned into training and testing datasets. Following the same approach as for vectors a structured approach was used to partition data into training and testing datasets, this was based on spatial autocorrelation of points based on Morans I, environmental similarity, and the number of points available to ensure even division of points between the training and testing datasets (flexsdm::*part_sband,* v1.3.4, Velazco et al. (2022)). (Roberts et al., 2017, Santini et al., 2021, Velazco et al., 2022). Individual species data were then combined for modelling into corvid (common raven, rook, carrion crow) or non-corvid (house sparrow, blackbird, barn swallow) representative groups. In the corvid group there were 33,132 (22,994 and 10,138 in the training and testing datasets respectively) presences and 32,002 (19,476 and 12,526 in the training and testing datasets respectively) background points. In the non-corvid group there were 43,085 (25,944 and 17,141 in the training and testing datasets respectively) presences and 42,273 (22,108 and 20,165 in the training and testing datasets respectively) background points.

Three individual correlative models were fitted using the *tune* functions in the *flexsdm* package, with selections using a threshold when the sum of sensitivity and specificity was highest (*max_sens_spec*). The correlative modelling approaches used were Random Forest (RF, flexsdm::*tune_raf,* v1.3.4, Velazco et al. (2022)), Generalized Boosted Regression (GBM, flexsdm::*tune_gbm,* v1.3.4, Velazco et al. (2022)), and Neural Network (NET, flexsdm::*tune_net, v1.3.4,* Velazco et al. (2022)). An Ensemble model based on the previous three individual models was also fit (*fit_ensemble, flexsdm)* (Velazco et al., 2022). Several performance metrics were calculated during model fitting and considered for model evaluation, these were True positive rate (TPR), True negative rate (TNR), Sorensen, Jaccard, F-measure (FPB), Omission rate (OR), True skill statistic (TSS), Area under curve (AUC), Boyce index and Inverse mean absolute error (IMAE) (Velazco et al., 2022, Valavi et al., 2022, Valavi et al., 2021, Hijmans and Elith, 2013, Roberts et al., 2017). These performance metrics have been described in detail in the vector section previously.

### Mechanistic species distribution models for vectors

To further validate our choice of species distribution models of vectors the mechanistic model CLIMEX was used, to ensure that areas predicted to be of higher suitability for *Culex* mosquitos using the correlative approaches were consistent with the known eco-physiological tolerance of *Cx. pipiens* and *Cx. modestus*. The mechanistic model was developed using the CLIMEX Compare Locations Genetic Algorithm (single species) function (v. 4) (Sutherst and Maywald, 1985, Kriticos et al., 2015). This was used to determine the most suitable parameter values based on the global presence datasets available alongside the literature and expert opinion to ensure the model was as reliable as possible (Kriticos et al., 2015, Sutherst and Maywald, 1985). Sensitivity analysis was carried out using the Compare Locations with Sensitivity Analysis function in CLIMEX, which showed that the most influential parameters for the ecoclimatic index calculation were soil moisture (*limiting low moisture, lower optimal moisture*), temperature (*lower optimal temperature)* and diapause (*diapause induction daylength, diapause termination temperature*) (Supplementary Information (SI) 3). As insufficient data was available to accurately parametrise for soil moisture constraints on both species the soil moisture index was removed from the model as incorrectly fitted parameters can greatly influence the suitability indices if the model is highly sensitive to them (Taylor and Kumar, 2012). The parameter values used in the CLIMEX model are shown in Table 2, and parameters not shown were not included in fitting the model. CLIMEX outputs were converted to a raster in ArcGIS Pro v. 3.0.1 (Esri, 2022) using the Point to Raster function, with the Ecoclimatic Index (EI) as the value field, and a cell size (i.e. resolution) of 0.167 degrees (approximately 1.85 x 1.85km). Validation of CLIMEX models has previously been carried out by using a presence dataset to determine how many known presence points fall within areas predicted to be suitable by the CLIMEX model (Taylor and Kumar, 2012). This approach of validation was used here, with environmental suitability categories created based on the ecoclimatic index with a score of 0 to 1 considered unsuitable, 1 to 10 as marginally suitable, 10 to 20 as favourable and >20 as very favourable. This followed the environmental classification approach used for the CLIMEX model for the mosquito *Aedes aegyptii* (Khormi and Kumar, 2014).

**Table 2:**
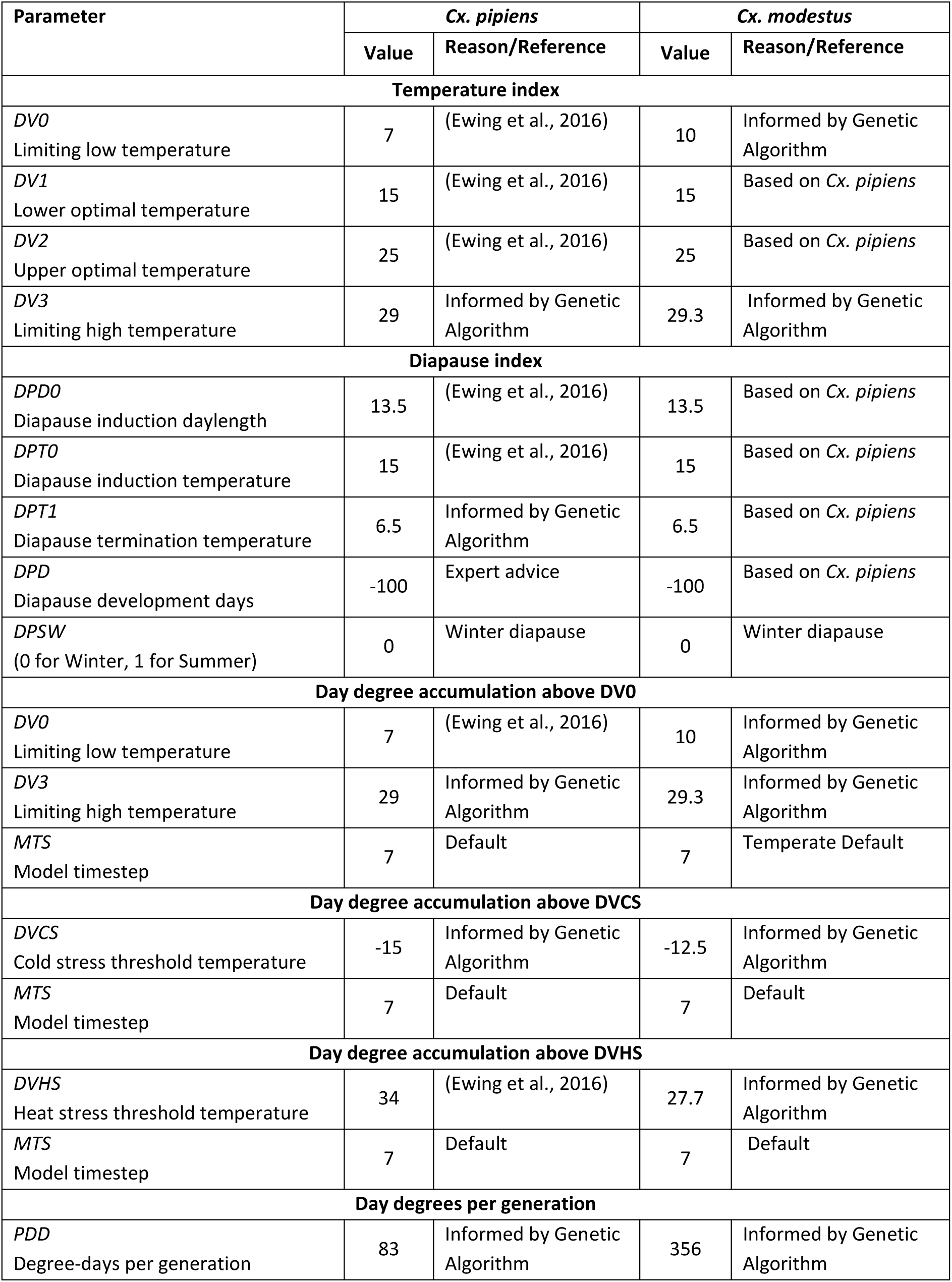
Parameter values used for CLIMEX modelling of *Cx. pipiens* and *Cx. modestus*.

#### Model predictions

Based on the global model, predictions of species distributions for the two *Culex* species across the UK were created (flexsdm:: *sdm_predict,* v1.3.4, Velazco et al. (2022)). For the MaxEnt binary predictions the threshold for suitability was based on the maximum sensitivity-specificity threshold (max_sens_spec), and for all other models continuous predictions were made.

To consider model transferability, predictions to regions outside the extent of the training data, were assessed using Mahalanobis distance and the shape method, with distance calculations based on the environmental distance between the training data (post-variable selection), and the area being projected into (flexsdm::*extra_eval,* v1.3.4, Velazco et al. (2022)) (Velazco et al., 2022, Velazco et al., 2023). The higher the Shape value, the greater the environmental distance is from the training data to the projection area (i.e., the more novel), indicating that areas with high Shape values have less reliable predictions than those with low Shape values (Velazco et al., 2023). Model transferability was measured both for the current and future environmental conditions under SSP1 and SSP5 for both *Cx. pipiens* and *Cx. modestus*. To further consider model extrapolation, partial dependence plots for the variables were created for each of the individual correlative models to determine how each model handled predictions to novel environmental conditions within the UK (flexsdm*::p_pdp*, v1.3.4, Velazco et al. (2022)).

Future predictions were made using the best performing model for *Cx. pipiens* and *Cx. modestus* considering the known distributions, the mechanistic CLIMEX model and the different performance scores. Future predictions were produced for the year 2100 based on the SSP126 and SSP585 (flexsdm:: *sdm_predict,* v1.3.4, Velazco et al. (2022)). Combined vector risk maps were then produced by multiplying the habitat suitability scores of cells for both *Cx. pipiens* and *Cx. modestus* for the current and future time periods to show overall vector risk based on habitat suitability up to the year 2100 for SSP1 and SSP5.

To produce a map of WNV risk the non-/corvid host species distribution maps were combined into one host risk map by summing the suitability score of all cells overlayed to give an overall host suitability score that included both representative corvid and non-corvid hosts. This host distribution model was then combined with the vector risk map by multiplying the host and vector maps together to produce an overall WNV risk map that considered both vectors and hosts. To further describe WNV risk in terms of human and equine health, this process was repeated to include layers representative of human population density both in the current environment and into the future for SSP1 and SPP5 in the year 2100 (Gao, 2020, Gao, 2017) and the current horse population density in the UK (Gilbert et al., 2022).

## Results

### Transferability of *Culex* species distribution models to areas and time-periods not represented by the presence data

Overall, the environmental conditions covered by the *Cx. pipiens* presence data represent those found across much of the UK, with only areas in northern Scotland (e.g., north-west Scotland, Shetland Islands) and very urban areas of England (e.g., London and Birmingham) showing as less represented (Figure 1a). Compared to *Cx. pipiens,* there was greater variation between the training data and predicted data for *Cx. modestus*, however many of the environmental conditions of the UK were covered by the data available, with London and north-west Scotland most poorly represented (Figure 1d). Similar patterns of environmental representation continued in future predictions for both species, with SSP5 showing more extrapolation was needed from the training data compared to SSP1 (Figure 1b-c for *Cx. pipiens* and Figure 1e-f for *Cx. modestus*). Partial dependence plots were created for each of the individual five correlative models to determine how each of the models’ handled data within the novel areas for both *Cx. pipiens* and *Cx. modestus* (Supplementary Information (SI) 4).

**Figure 1a-f:**
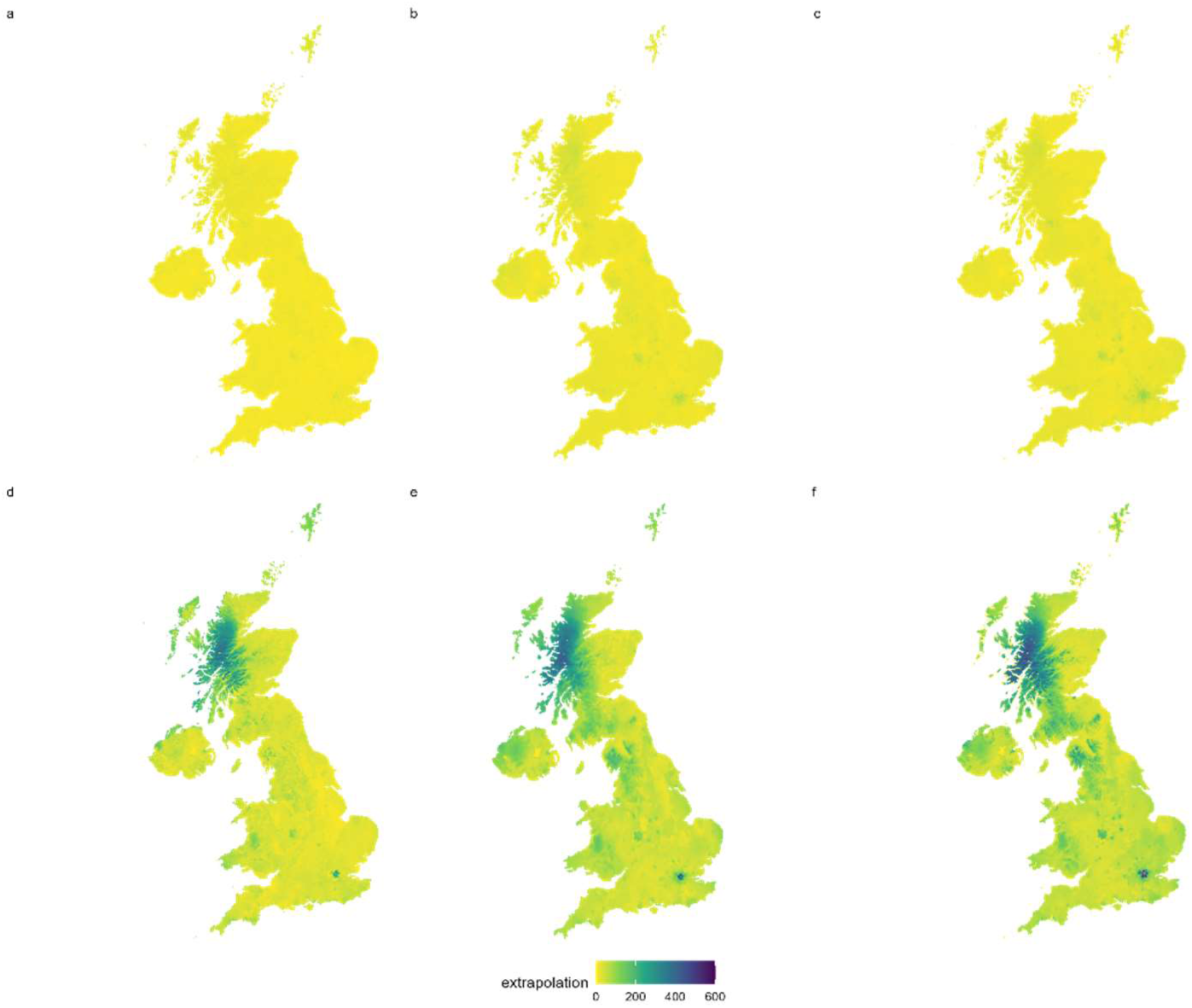
The Shape values for extrapolation of the study area based on the presence data available for the *Culex* species distribution models. The lower the shape value the more similar the regions are, thus the higher values represent environmental conditions poorly represented by the presence data available. Extrapolation values are shown for *Culex pipiens* for the a) current and b) future (year 2100) environment under scenario SSP1 and c) SSP5, and for *Culex modestus* for the d) current and e) future (year 2100) environment under scenario SSP1 and f) and SSP5.

### Correlative species distribution models for vectors in the current climate and environmental conditions

#### Culex pipiens

The overall pattern of the individual *Cx. pipiens* distribution models was similar across the UK, with areas of higher suitability around the north-east Scotland, south-east, central, and north-west in England and coastal Wales (Figure 2a-f). In all models the areas with lowest suitability for *Cx. pipiens* were middle-east England and northern Scotland. However, it was the areas of lower suitability that had higher variability with GBM and SVM models predicting wider spread unsuitability compared to the RF, MaxEnt and Net models (Figure 2a-f).

**Figure 2a-f:**
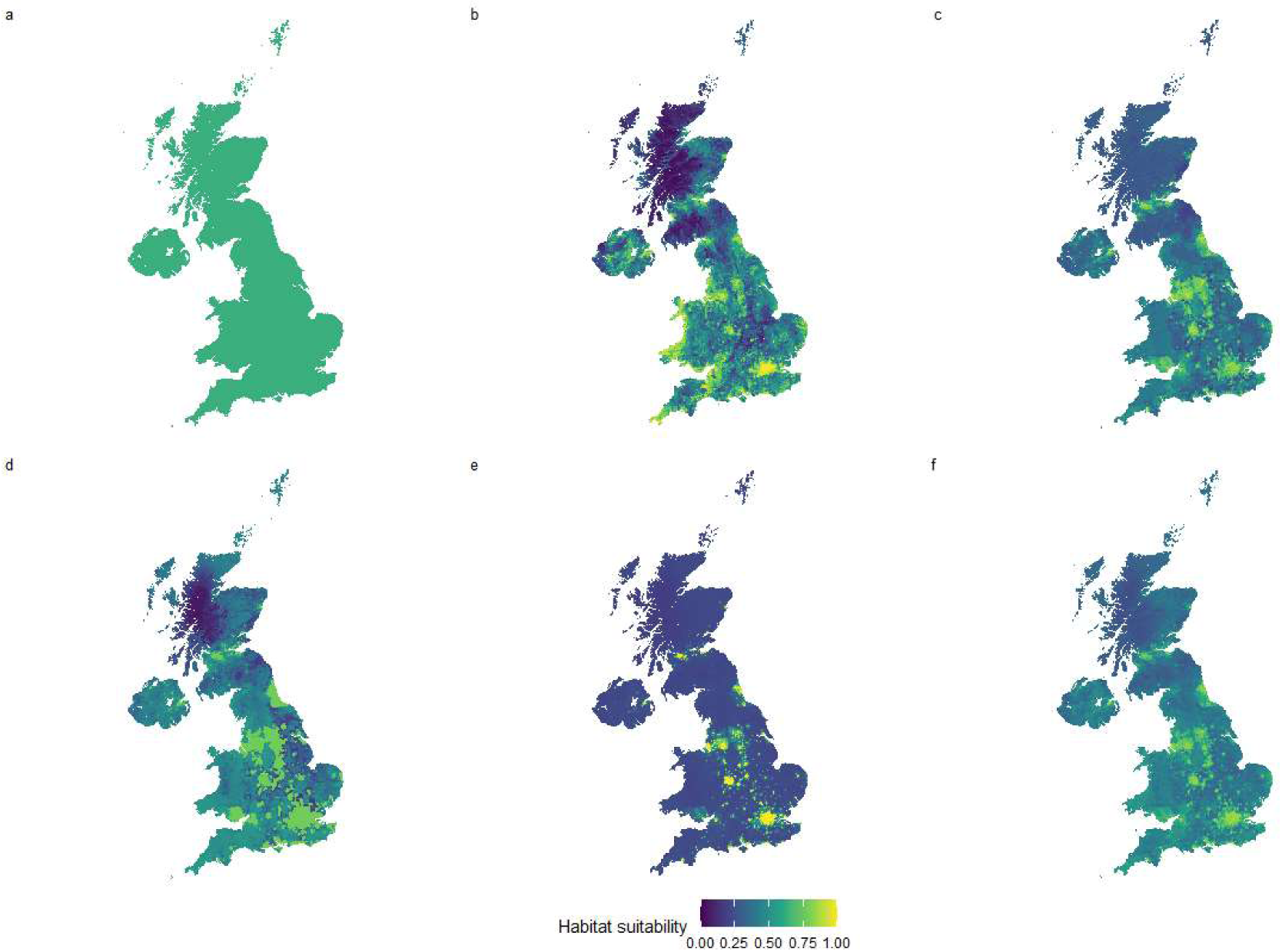
Species distribution models for *Culex pipiens* using a) MaxEnt, b) Random Forest, c) Generalized Boosted Regression, d) Neural Network, e) Support Vector Machine and f) A mean weighted Ensemble model of a – e, that was weighted based on model performance.

Model performance can be measured in many ways and, based on the different performance metrics, the most consistently high scoring model for *Cx. pipiens* was the Ensemble model, with the Sorensen index, Jaccard, FPB, TSS and AUC all being the highest, with the remaining scores being highest for either the GBM (TNR, OR, IMAE) or RF (TPR, Boyce) models. Considering this, the *Cx. pipiens* model that performed best when predicting habitat suitability in the UK was the mean weighted Ensemble model (Table 3). This model reflects the known distribution of *Cx. pipiens* in the UK, with higher suitability in the south compared to the north (European Centre for Disease Prevention and Control and European Food Safety Authority, 2023), and that predicted by the CLIMEX model which is discussed further in the mechanistic species distribution model section.

**Table 3:** Model performance scores and standard deviation (sd) of the Culex models. The performance metrics are True positive rate (TPR), True negative rate (TNR), Sorensen, Jaccard, F-measure (FPB), Omission rate (OR), True skill statistic (TSS), Area under curve (AUC), Boyce index and Inverse mean absolute error (IMAE). Higher values indicate better model performance.

|  |  | TPR |  | TNR |  | Sorensen |  | Jaccard |  | FPB |  | OR |  | TSS |  | AUC |  | Boyce |  | IMAE |  |
| --- | --- | --- | --- | --- | --- | --- | --- | --- | --- | --- | --- | --- | --- | --- | --- | --- | --- | --- | --- | --- | --- |
|  | score<br>range | 0 to 1 |  | 0 to 1 |  | 0 to 1 |  | 0 to 1 |  | 0 to 2 |  | 0 to 1 |  | -1 to 1 |  | 0 to 1 |  | -1 to 1 |  | 0 to 1 |  |
|  |  | mean | sd | mean | sd | mean | sd | mean | sd | mean | sd | mean | sd | mean | sd | mean | sd | mean | sd | mean | sd |
| Culex pipiens |  |  |  |  |  |  |  |  |  |  |  |  |  |  |  |  |  |  |  |  |  |
| SVM |  | 0.70 | 0.10 | 0.71 | 0.05 | 0.50 | 0.21 | 0.35 | 0.19 | 0.70 | 0.38 | 0.30 | 0.10 | 0.42 | 0.14 | 0.75 | 0.10 | 0.70 | 0.18 | 0.69 | 0.01 |
| Neural Network |  | 0.63 | 0.19 | 0.76 | 0.09 | 0.51 | 0.16 | 0.35 | 0.15 | 0.69 | 0.30 | 0.37 | 0.19 | 0.39 | 0.10 | 0.73 | 0.08 | 0.87 | 0.04 | 0.71 | 0.01 |
| GBM |  | 0.60 | 0.20 | 0.80 | 0.06 | 0.51 | 0.19 | 0.35 | 0.17 | 0.71 | 0.34 | 0.40 | 0.20 | 0.40 | 0.14 | 0.75 | 0.09 | 0.79 | 0.24 | 0.71 | 0.03 |
| Random Forest |  | 0.73 | 0.05 | 0.67 | 0.16 | 0.49 | 0.21 | 0.34 | 0.18 | 0.67 | 0.37 | 0.27 | 0.05 | 0.40 | 0.12 | 0.74 | 0.06 | 0.98 | 0.03 | 0.69 | 0.01 |
| MaxEnt |  | 0.72 | 0.08 | 0.67 | 0.03 | 0.49 | 0.16 | 0.33 | 0.14 | 0.67 | 0.28 | 0.28 | 0.08 | 0.40 | 0.05 | 0.74 | 0.06 | 0.96 | 0.06 | 0.60 | 0.00 |
| Ensemble mean weighted |  | 0.65 | 0.21 | 0.77 | 0.10 | 0.52 | 0.17 | 0.36 | 0.15 | 0.72 | 0.30 | 0.35 | 0.21 | 0.42 | 0.11 | 0.76 | 0.08 | 0.95 | 0.02 | 0.68 | 0.01 |
| Culex modestus |  |  |  |  |  |  |  |  |  |  |  |  |  |  |  |  |  |  |  |  |  |
| SVM |  | 0.68 | 0.12 | 0.66 | 0.11 | 0.5 | 0.02 | 0.33 | 0.02 | 0.67 | 0.04 | 0.32 | 0.12 | 0.34 | 0.01 | 0.66 | 0.02 | 0.89 | 0.07 | 0.64 | 0.02 |
| Neural Network |  | 0.79 | 0.17 | 0.61 | 0.16 | 0.53 | 0.03 | 0.36 | 0.03 | 0.72 | 0.06 | 0.21 | 0.17 | 0.39 | 0.01 | 0.68 | 0.08 | 0.93 | 0.08 | 0.67 | 0.02 |
| GBM |  | 0.58 | 0.19 | 0.76 | 0.16 | 0.5 | 0 | 0.33 | 0 | 0.67 | 0 | 0.42 | 0.19 | 0.34 | 0.03 | 0.63 | 0.03 | 0.95 | 0.06 | 0.71 | 0.04 |
| Random Forest |  | 0.32 | 0.18 | 0.93 | 0.04 | 0.4 | 0.17 | 0.26 | 0.13 | 0.52 | 0.26 | 0.68 | 0.18 | 0.25 | 0.14 | 0.56 | 0.08 | 0.72 | 0.15 | 0.67 | 0 |
| MaxEnt |  | 0.63 | 0.21 | 0.7 | 0.09 | 0.5 | 0.1 | 0.33 | 0.09 | 0.67 | 0.18 | 0.37 | 0.21 | 0.33 | 0.12 | 0.61 | 0.08 | 0.73 | 0.05 | 0.54 | 0.1 |
| Ensemble mean threshold |  | 0.66 | 0.27 | 0.7 | 0.18 | 0.52 | 0.07 | 0.35 | 0.07 | 0.7 | 0.13 | 0.34 | 0.27 | 0.36 | 0.08 | 0.67 | 0.05 | 0.96 | 0.02 | 0.72 | 0.05 |

#### Culex modestus

Overall, the pattern of suitability across the UK for *Cx. modestus* was relatively similar for the RF, GBM and Net models, showing predominantly unsuitable habitat, apart from areas of southern and central England, particularly in the south-east (Figure 3a-f). The MaxEnt and SVM models showed widespread suitability across the UK, though most of these areas were scored as relatively low suitability compared to the other models (Figure 3a-f).

**Figure 3a-f:**
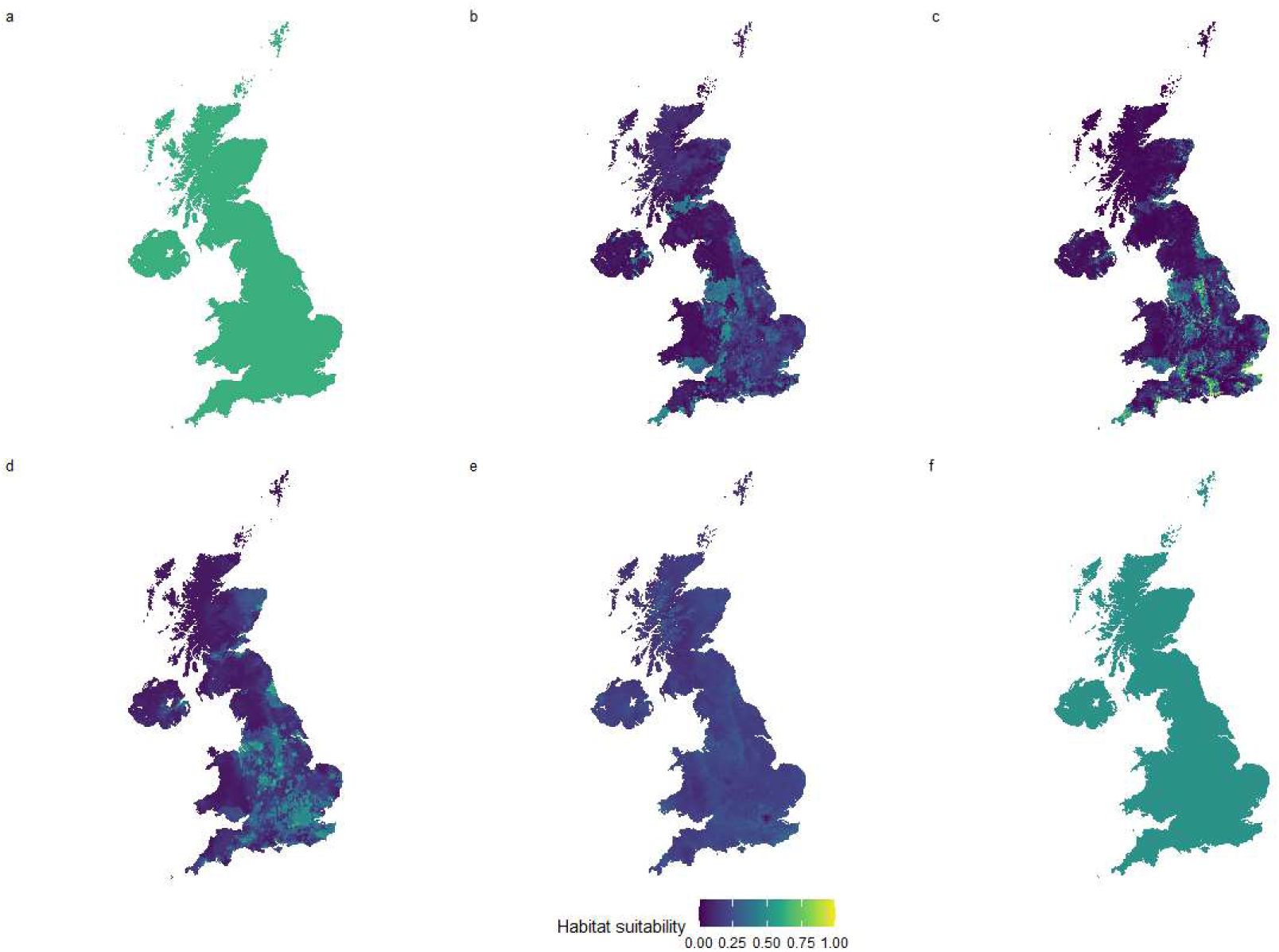
Species distribution models for *Culex modestus* using a) MaxEnt, b) Random Forest, c) Generalized Boosted Regression, d) Neural Network, e) Support Vector Machine and f) A mean threshold Ensemble model of a – e based on cells with suitability above the maximum sensitivity and specificity threshold.

The best performing model for *Cx. modestus* was the Net model, which scored highest in the TPR, Sorensen, Jaccard, FPB, TSS and AUC performance measures. Both the RF and Ensemble Mean Threshold models also scored highly for certain metrics (TNR/OR and Boyce/IMAE, respectively).

Considering this, the Net model for *Cx. modestus* was taken to be the most reliable prediction of habitat suitability in the UK (Table 3). The pattern of suitability for the Net model aligns with the known presence data for this species, which is predominantly associated with the North Kent marshes in the south-east in the UK (Golding et al., 2012) and with that predicted by the CLIMEX model which is discussed in the mechanistic species distribution model section.

### Mechanistic species distribution models for vectors in the current climate and environmental conditions

#### Culex pipiens

Based on the CLIMEX model, Southern England is most suitable for *Cx. pipiens,* with a higher ecoclimatic index compared to central and northern regions. Overall, much of the UK is suitable for *Cx. pipiens* as an ecoclimatic index greater than 1 indicates that some growth could occur. The only region calculated to be unsuitable is an area of higher elevation in central-north Scotland (Figure 4a). Validation of the CLIMEX model to known presences was able to use 655 *Cx. pipiens* presence points that fell within the study area, and these were assigned to categories based on the ecoclimatic index score of the model. Of these, most presence points fell into areas considered to be very favourable (N = 646), with the remaining presences located in favourable (N = 8) or marginally suitable (N = 1) areas. No presences were recorded in areas considered unsuitable based on the ecoclimatic index.

**Figure 4a,b:**
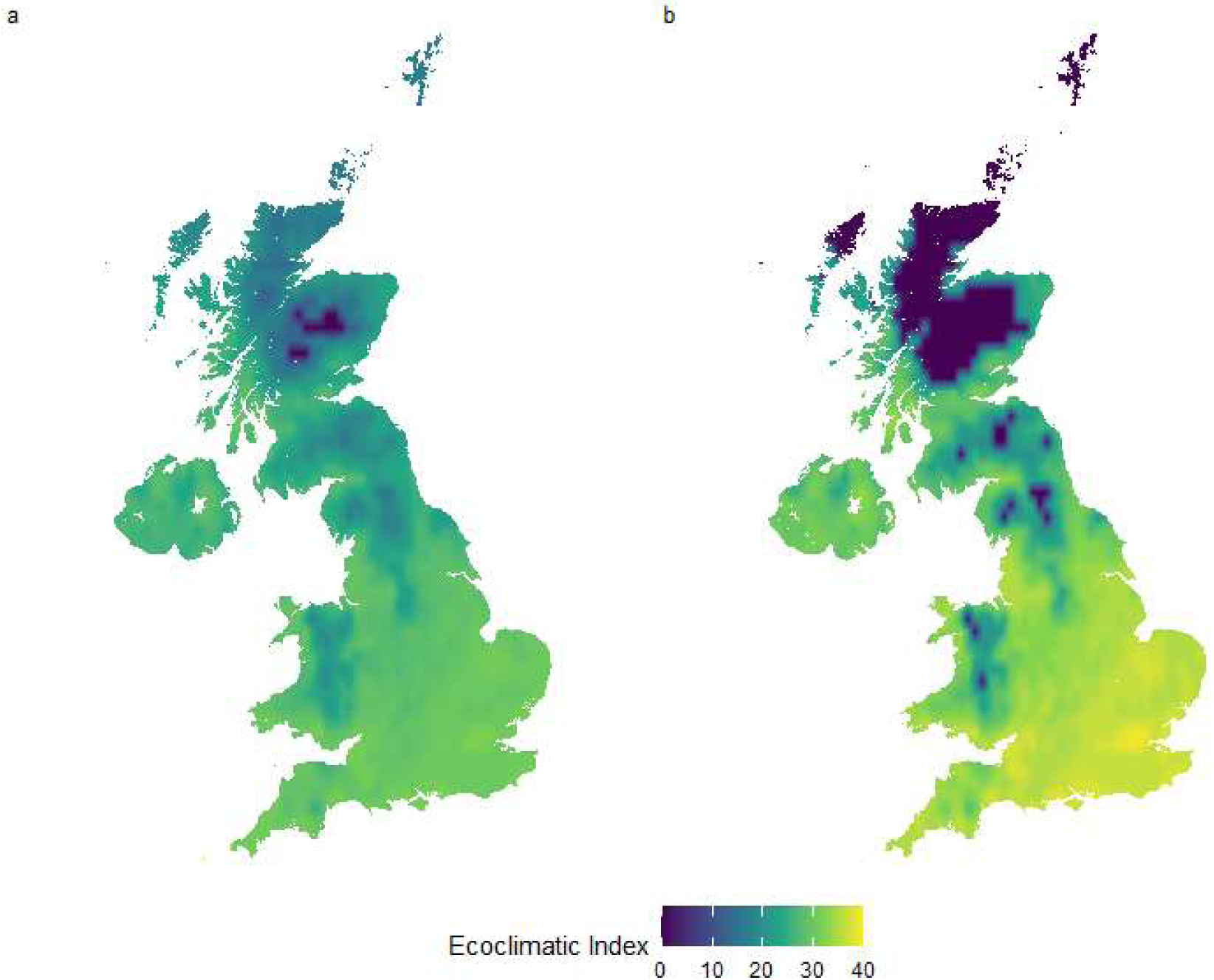
The ecoclimatic index of the UK for a) *Cx. pipiens* and b) *Cx. modestus*. An ecoclimatic index greater than 1 indicates some survival can occur, with the higher the number the more suitable the area is climatically for the species.

#### Culex modestus

The CLIMEX model for *Cx. modestus* predicted that the south-east of England had the highest ecoclimatic index in the UK, whereas much of northern and central Scotland was unsuitable (ecoclimatic index was very low or 0 suggesting that no population growth could occur; Figure 4b).

Validation using known presence records showed that all 6 *Cx. modestus* records for the UK were in areas that were very favourable based on the ecoclimatic index, however, the limited sample size does reduce the reliability of validation for this species. Therefore, as has previously been done when modelling with CLIMEX (Early et al., 2022), visual comparisons to known and predicted *Cx. modestus* distributions were also considered and these did align with the output of the CLIMEX model (Golding et al., 2012, European Centre for Disease Prevention and Control, 2019).

### West Nile Virus case study; current and future predictions of risk in the UK

#### Risk based on *Culex* vector distribution

Considering future projections for both *Cx. pipiens* and *Cx. modestus*, there are many areas across the UK that see an increase or decrease in suitability for both SSP1 and SSP5 scenarios. Increases in suitability are predominantly in areas currently identified as highly suitable, such as across much of eastern England (Figure 5b-e). In the year 2100, the overall pattern of change for the SSP5 scenario is very similar to that for SSP1, however the changes are predicted to be greater, reflecting the increased change in the environment with the SSP5 scenario (Figure 5b-e).

**Figure 5a-e:**
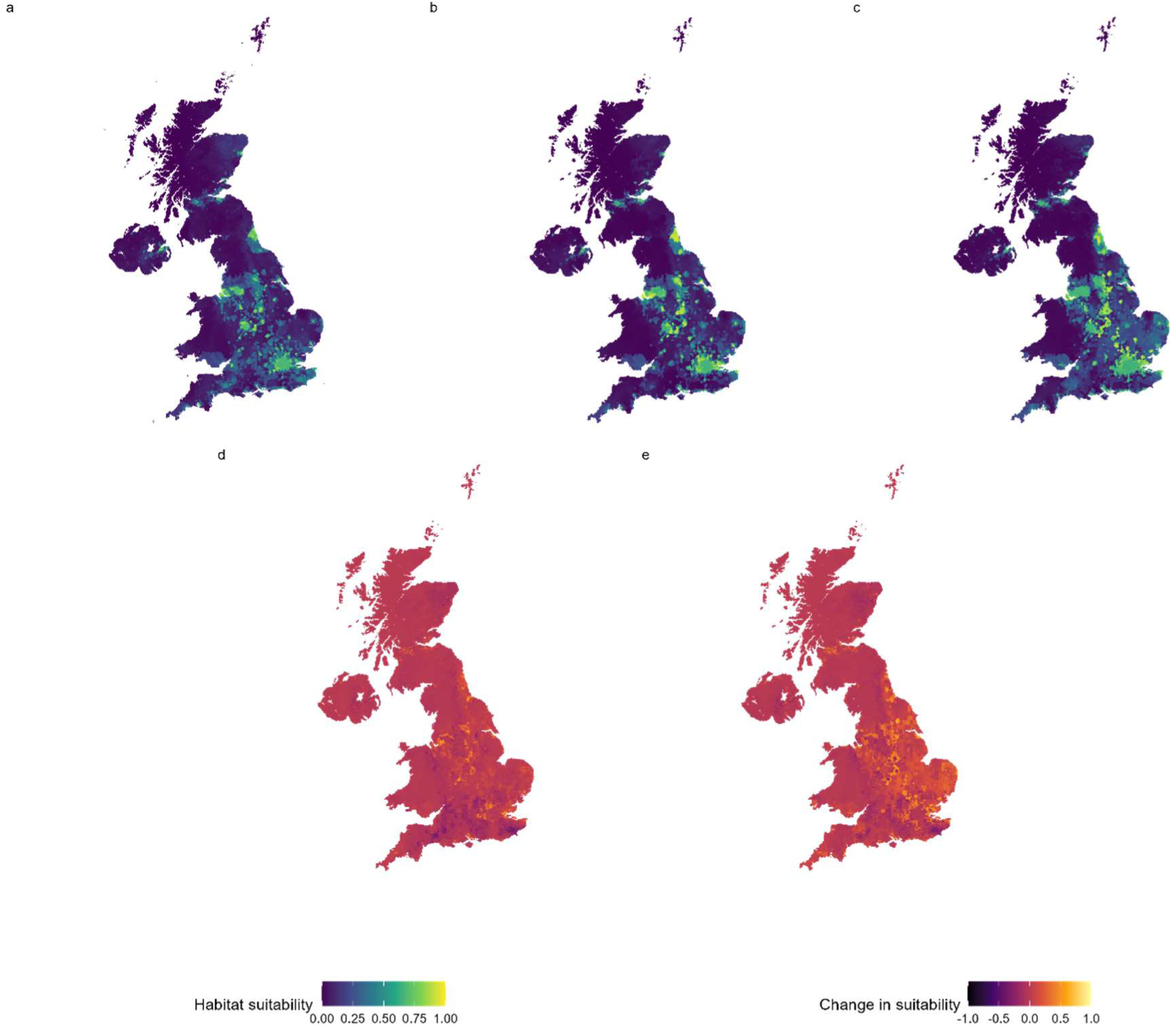
Habitat suitability for *Culex spp.* (*Cx. pipiens* and *Cx. modestus*) in a) the current climate and in the year 2100 under b) SSP1 and c) SSP5 scenarios, and the change in suitability between the current and future climates for scenario d) SPP1 and e) SSP5.

#### Risk based on *Culex* and representative avian host distribution

To get an improved understanding of WNV risk, a species distribution map of the representative corvid and non-corvid hosts was also created (Figure 6a-c). The results here focus on the species distribution modelling of vectors as the primary focus of this study; for full results of the species distribution models for the avian hosts see Supplementary Information (SI) 5. Similar to the vector distribution predictions, when considering the future environment some areas showed an increase in suitability in south-east England and north-west Scotland whereas others had reduced suitability such as in the north-east and central England. Overall there was much more widespread suitability across the UK for host species, including areas of high suitability in northern Scotland which was largely unsuitable for both vectors (Figure 5a-e, Figure 6a-c).

**Figure 6a-c:**
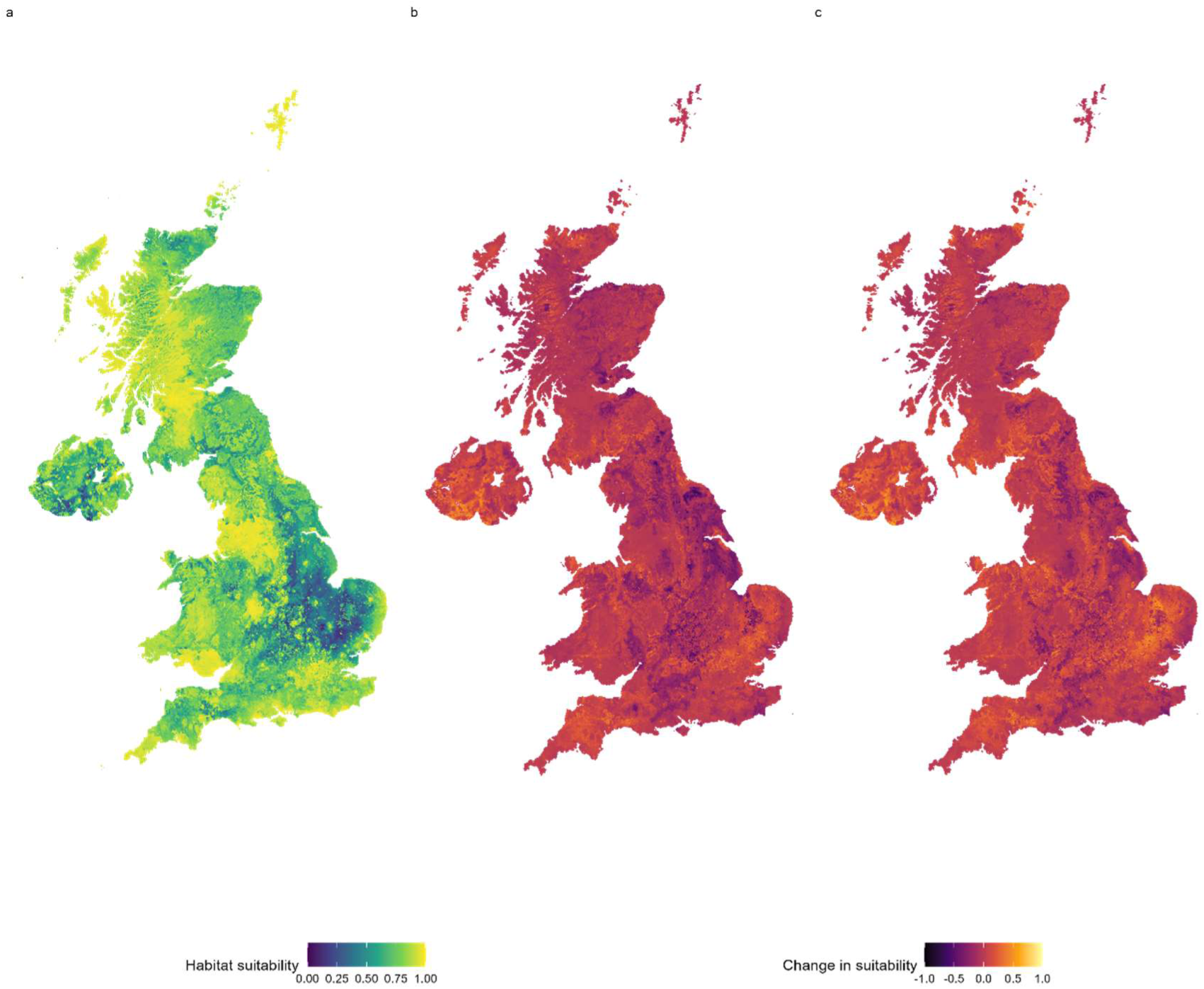
Habitat suitability for the combined, representative host species (corvid and non-corvid) in a) the current climate and the change in suitability by year 2100 under scenario b) SSP1 and c) SSP5. For full details of predictions for host suitability see Supplementary Information (SI) 5.

By combining vector and host distribution maps a WNV risk map was produced to show areas of high suitability for both vectors and hosts, as these areas are likely to have increased viral transmission and a higher likelihood of the virus establishing. The WNV risk map showed that the greatest risk is in England, particularly urbanised areas in the south-east, north-east and north-west (Figure 7). Predicting into 2100 revealed that many of the areas currently deemed to be at risk will have an increased risk under both SSP1 and SSP5 scenarios and, as expected based on the host and vector maps, this change will be greater under SSP5 (Figure 7).

**Figure 7a-e:**
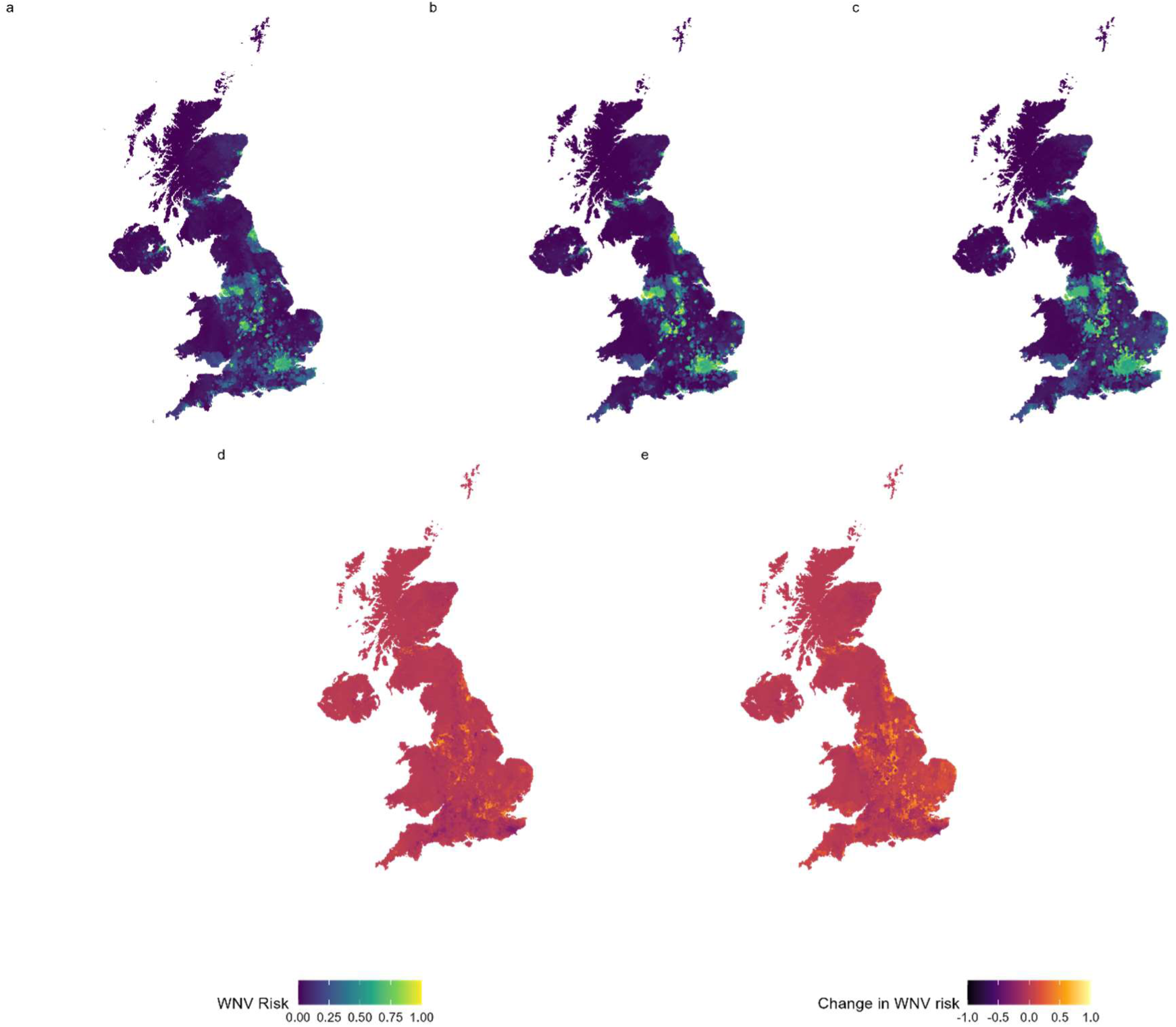
The WNV risk in the UK calculated by considering both vector (*Cx. pipiens*, *Cx. modestus*) and host (corvid and non-corvid) species distribution maps to infer overall risk. The figure shows a) the current risk, b) predicted risk in 2100 under scenario SSP1, c) the predicted risk in 2100 under scenario SSP5, and the change in risk by 2100 under scenario d) SSP1 and e) SSP5.

#### Risk based on *Culex* vectors, representative avian hosts and non-target human and equine hosts

To further define risk in terms of human and equine health, risk maps based on vector and host distributions were overlaid with maps reflecting human and horse densities in the UK. As expected, the greatest risk to horses and humans coincided with areas of high suitability for vector and host populations, such as the south-east and north-west of England (Figure 8a-d). Only human density estimates are available for the future, and these revealed a very similar pattern of risk under both SSP1 and SSP5, with at-risk areas expanding following the predicted spread of urbanised areas with higher population densities (Figure 8a-d).

**Figure 8a-d:**
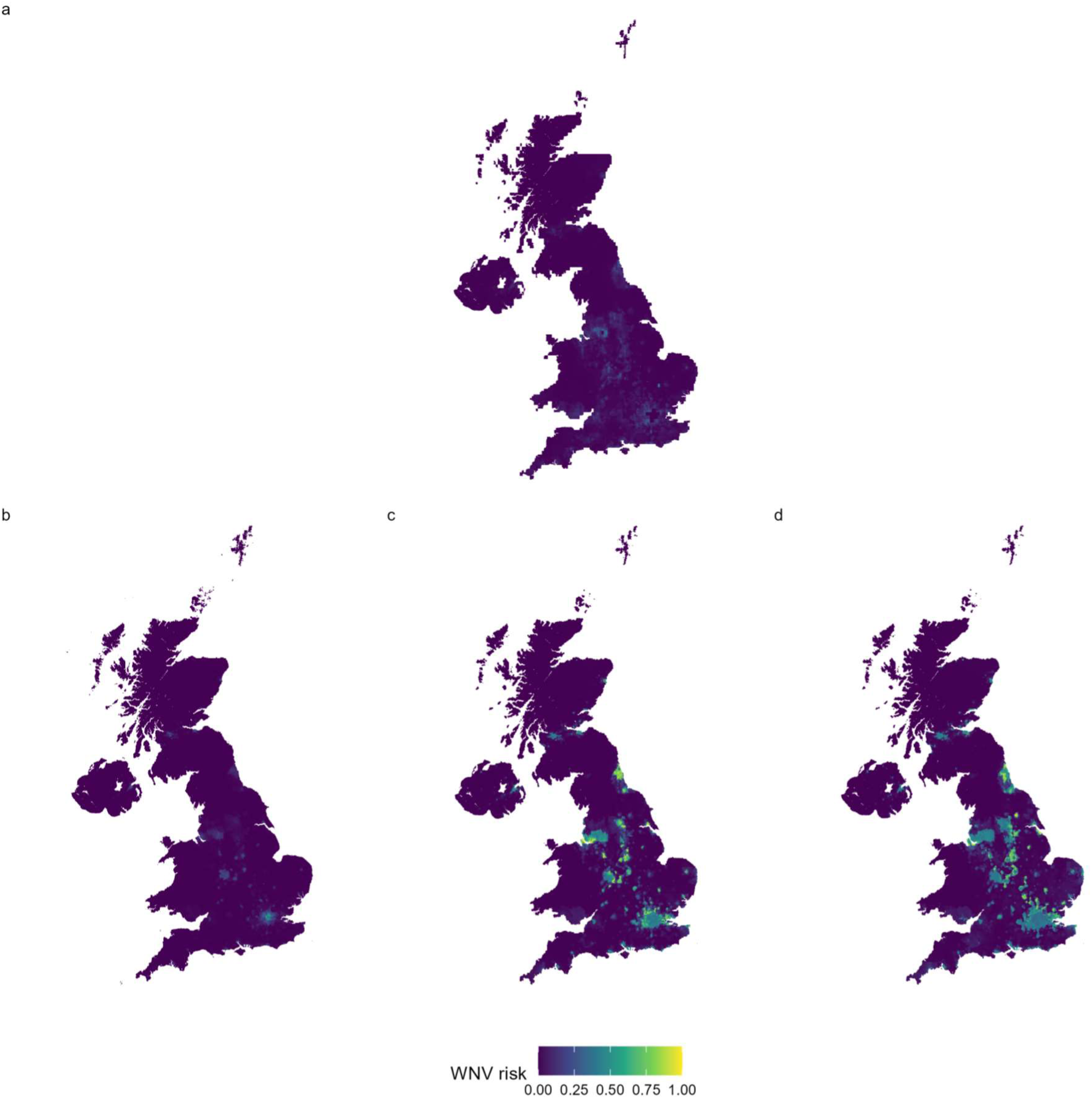
The WNV risk in the UK calculated by considering both vector (*Cx. pipiens*, *Cx. modestus*), target reservoir host (corvid and non-corvid) and dead-end host species distribution maps to infer overall risk. The figure shows a) the risk to UK horses in the current climate, b) the risk to the human population in the current climate, c) the risk to the human population in 2100 under scenario SSP1 and, D) the risk to the human population in 2100 under scenario SSP5.

## Discussion

The focus of this study was to determine a general framework of using species distribution modelling of vectors and hosts to highlight areas at risk of vector-borne diseases in the UK. Here, it is proposed that correlative species distribution models can be used to predict vector distributions, and that mechanistic modelling approaches can help to validate correlative models. These vector species distribution models can then be merged with correlative target host species distribution models to assess risk from disease, with further refinements of disease risk predictions by then combining with data for dead-end hosts such as humans, equines, or livestock.

### Implications for disease risk in the UK

Using this approach, a case study on WNV was carried out and a risk map produced, showing that hotspots of WNV risk exist across the UK, with the greatest risk being in the south-east and north-west England. When projecting this risk into the future under SSP1 and SSP5 environmental scenarios some areas have an increased risk, including much of eastern England and hotspots in the north-west. This risk was also considered in relation to both human and horse population densities and, as expected, this showed disease risk hotspots located in densely populated areas where both vectors and hosts were likely to be present. To further improve the risk map for horses, improved horse population density estimate maps for the UK will be required as this is currently an area with limited data availability, with current predictions likely to be underestimated. However, the current map is likely to reflect the overall distribution of horses and thus show the potential risk to horses within the UK. As such, these risk maps will be important for targeting surveillance so any disease incursion or outbreak can be detected quickly, enabling a rapid response that aims to control the disease prior to spread and establishment.

Whilst this study has focussed on WNV, there are also other pathogens spread by *Cx. pipiens* and *Cx. modestus*, such as Usutu virus, avian malaria (*Plasmodium spp.*) and heartworm (*Difilaria immitis*) (Soto and Delang, 2023, Farajollahi et al., 2011). The vector risk maps presented in this study will also represent areas of higher risk from these diseases in the UK and could be refined further using the approach described here to combine vector and relevant non-/target host species distribution models. Using this approach could improve our understanding of the risk of diseases spread by *Culex* vectors across the UK. Furthermore, this methodology could also be tailored for other vectors, countries or regions to determine disease risk based on the likelihood of vector presence elsewhere.

### Correlative and mechanistic approaches to species distribution modelling of *Culex* vectors in the UK

Whilst avian species distribution modelling was carried out to represent WNV host availability across the UK, carefully selected species were used to create representative host models. This is due to *Culex* species not showing a strong host preference for specific avian species and many different bird species being competent hosts for WNV (Brugman et al., 2013, Brugman et al., 2017, Centers for Disease Control and Prevention, 2021, Ziegler et al., 2022, Bergmann et al., 2023, Bessell et al., 2016). Therefore, vectors may be a key limiting factor in the spread of WNV and have higher habitat specificity so were the primary focus on this study and much of this discussion will focus on the species distribution modelling of the vectors.

The species distribution models of *Cx. pipiens* and *Cx. modestus* highlighted that both species had areas of high suitability across the UK, with these areas often overlapping, highlighting similarities in the species environmental needs. The transferability of the *Culex* data used to train the models is important to consider as the WNV risk models are predominantly driven by vector habitat suitability both in the current and future maps. Here, reviewing extrapolation of presence data with the Shape method proposed by Velazco *et al*. (2023) showed that the ability to predict to current and future climate in the UK was good for *Cx. pipiens* but more limited for *Cx. modestus*. This is to be expected as *Cx. modestus* is an invasive species in the UK that may not have reached its full range, and data availability is much more limited for this species (Golding et al., 2012, Soto and Delang, 2023, Elith et al., 2010, Venette, 2017). This further highlights the importance in improving current knowledge of vector distributions, especially for species that have very limited data, such as *Cx. modestus*.

Selecting which species distribution models are most appropriate can be a challenge, as each model can produce markedly different results and cross-validation scoring approaches can vary considerably as well as being strongly affected by biases in the data (Elith et al., 2006, Elith et al., 2010, Hijmans and Elith, 2013, Roberts et al., 2017, Valavi et al., 2022). Throughout the modelling process used in this study, steps were taken to ensure that biases were minimised. For example, some performance scores can be artificially inflated by how data is partitioned into testing and training datasets for cross-validation (Velazco et al., 2022, Roberts et al., 2017). To overcome this host and vector data were partitioned based on environmental conditions in the model calibration area. This was particularly important to consider for vector data as it is more limited than the avian host data, with strong spatial biases in presence locations due to insect sampling methods typically relying on trapping which can have biases due to trap attractiveness and locations (Koch, 2021, Petrovskii et al., 2012).

The model that performed best was taken forward for both vectors and hosts. For *Cx. pipiens* this was the mean-weighted Ensemble model whereas for *Cx. modestus* this was the Neural Network model, highlighting how model performance can vary for different species. Species-specific variations in model performance have been recorded previously for mosquito species in Europe (European Centre for Disease Prevention and Control, 2019), as well as across 225 other species (Valavi et al., 2022). These differences in model suitability and scoring can be due to a number of reasons, such as variation in sampling effort, balances between presence and absences and species characteristics (e.g., species niches) (Valavi et al., 2022). Whilst performance metrics were used in this study, visual comparisons to known distributions, previous predictions, and to the CLIMEX mechanistic model were also considered to aid model selection (European Centre for Disease Prevention and Control, 2019, European Centre for Disease Prevention and Control and European Food Safety Authority, 2023, Golding et al., 2012). This was important as it allowed consideration of biological limits which can reflect geographical survival ranges of vector species.

There was more data available on *Cx. pipiens* to parametrise the CLIMEX model, and the genetic algorithm was used to further inform some parameters based on known presence data (Ewing et al., 2016, Ewing et al., 2017, Ewing et al., 2019). By comparing the CLIMEX output to presence data available to inform on model performance, most presences were in areas considered to be highly suitable for *Cx. pipiens* and no presences were in areas predicted to be unsuitable, indicating the CLIMEX model performed well. Parameterising the CLIMEX model for *Cx. modestus* was more challenging than for *Cx. pipiens* as data was much more limited (Soto and Delang, 2023). Moreover, *Cx. modestus* has more specific habitat requirements for different life-stages, with larvae typically recorded in permanent water bodies only (Soto and Delang, 2023). As such, land cover is not considered in the CLIMEX modelling approach. Modelling using CLIMEX for species with limited distributions has been carried out based on the upper and lower limits of the observed distribution (Khormi and Kumar, 2014), or by matching to the known distribution of a species (Poutsma et al., 2008, Venette, 2017). However, for invasive species with limited distributions and poor data availability these approaches could lead to errors in estimating species distribution, as both rely on absences in distribution data being true absences and may not reflect the biological limits of a species (Venette, 2017). Therefore, to overcome these challenges the *Cx. modestus* CLIMEX model relies on the genetic algorithm results, supplementing missing parameters with those used for *Cx. pipiens* as it is a closely related species that also has winter diapause (Soto and Delang, 2023). The resulting *Cx. modestus* model performed well in model validation, with all presence data available in areas considered to be very favourable and visual comparisons showing alignment with the known distribution of this species (Golding et al., 2012, European Centre for Disease Prevention and Control, 2019). This would suggest that the model is informative, however, to further improve the CLIMEX model, extensive research to refine model parameterisation would be beneficial for *Cx. modestus*, especially those parameters that show high levels of sensitivity (i.e. temperature limits, soil moisture), which can lead to strong variations in suitability predictions based on ecoclimatic indices (Taylor and Kumar, 2012).

Comparisons of the mechanistic and correlative models showed similarities for *Cx. modestus*, with both approaches predicting higher suitability in the south-east and very low suitability in northern-Scotland. Similarly, the *Cx. pipiens* mechanistic model closely reflects the output of the correlative models, with both showing southern England and the western coastal areas of Wales as more suitable than more northern areas. The CLIMEX model does not highlight urban areas as more suitable for *Cx. pipiens* however the correlative models can identify this, which is expected as *Cx. pipiens* is commonly associated with urbanisation (Ruybal et al., 2016, Krol et al., 2024, Townroe and Callaghan, 2014). This is likely due to land use being included in the correlative models whereas the CLIMEX model is based solely on biological limits and climate information.

Therefore, for vectors with strong associations to land use types correlative models may be more suitable, with CLIMEX used to support where the climatic conditions are suitable for vector survival and reproduction, with correlation between the two modelling approaches indicative of strong models.

## Conclusion

By combining both vector and host distributions it is possible to assess potential risks from novel diseases in the UK. This can highlight where a pathogen is most likely to establish due to increased environmental suitability for both vectors and hosts, both presently and in the future. This study has highlighted that the risk of WNV risk establishment is expected to increase in some areas whilst decreasing in others, leading to areas of high and low risk across the UK in the future. Overall, a novel method, combining correlative and mechanistic vector species distribution models, correlative host species distribution models and non-target host data to consider disease risk is presented. It is hoped that this approach will be transferable to other diseases and geographic regions in the future.

## • data availability statement

All data underlying this work are held in publicly accessible repositories and can be downloaded for use in research under licence. Outputs can be made available under license by contacting the corresponding author.

## • funding statement

This work is funded by EXSE0566: Development of improved tools and approaches for control of zoonotic arthropod-borne diseases of animals and their vectors Funded by the Department for Environment, Food and Rural Affairs and the Scottish and Welsh Governments and was co-funded by the European Union’s Horizon Europe Project 101136346 EUPAHW.

## • conflict of interest disclosure

The authors declare no conflicts of interest.

## • ethics approval statement

n/a

## • patient consent statement

n/a

## • permission to reproduce material from other sources

n/a

## • clinical trial registration

n/a

## Notes

### Competing Interest Statement

The authors have declared no competing interest.

### Summary of Updates

The funding statement has been updated.

