## Supplementary Information (SI) for "Using correlative and mechanistic species distribution models to predict vector-borne disease risk for the current and future environmental and climatic change: a case study of West Nile Virus in the UK"

### SI.1: Correlation of predictor variables

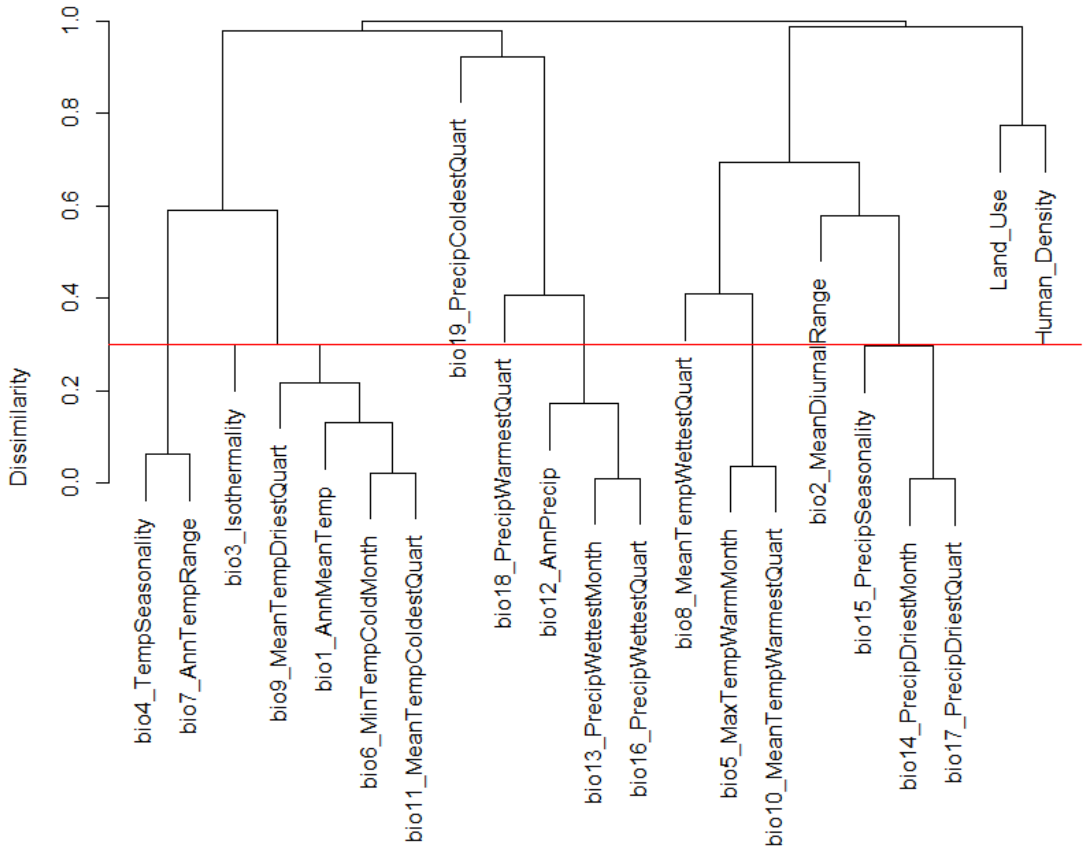


*Figure S. 1: A dendrogram showing correlation clusters of predictor variables used in this study. Pearson correlation coefficient was used to remove climatic variables with a dissimilarity index of <0.3 as this indicated a correlation coefficient >0.7, as represented by the red line. Following removal of correlated variables, the remaining predictor variables for species distribution modelling were bio2 (mean diurnal range), bio8 (mean temperature of the wettest quarter), bio9 (mean temperature of the driest quarter), bio14 (precipitation of the driest month), bio15 (precipitation seasonality), bio18 (precipitation of the warmest quarter), bio19 (precipitation of the coldest quarter), land use and human population density.*

### SI. 2: Calibration area determination for vector correlative species distribution modelling

To determine the most suitable calibration area to use for species distribution modelling of vectors, comparisons were made between calibration areas of 5km, 50km and 100km around presence points for both *Cx. pipiens* and *Cx. modestus*. Calibration areas were created using the *calib_area* function in the *flexsdm* package (v1.3.4, Velazco et al. (2022)). Due to variations in the size of the calibration area, the number of background points that could be created varied due to being unable to find a sufficient number of empty cells to place background points in when using the *randomPoints* function (*dismo*, v1.3-14, Hijmans et al. (2023)). For *Cx. pipiens* the target number of background points was 4,577 (3x number of presences) however the 5km calibration area had 920, the 50km calibration area had 2,484 and the 100km calibration had 4,577. This variation in the number of background points also occurred for *Cx. modestus* where the target number was 141 which was achieved by the 100km calibration area and almost achieved by the 50km calibration area which had 140 background points, however, the 5km calibration area only had 67 background points. Pearson correlation coefficient was used to remove variables with a correlation coefficient >0.7. Following this, the remaining variables included were bio2 (mean diurnal range), bio8 (mean temperature of the wettest quarter), bio9 (mean temperature of the driest quarter), bio14 (precipitation of the driest month), bio15 (precipitation seasonality), bio18 (precipitation of the warmest quarter), bio19 (precipitation of the coldest quarter), land use and human population density. A range of modelling approaches and different performance metrics were considered when selecting which calibration area and model to choose, and these are shown in Table S. 2. For both *Cx. pipiens* and *Cx. modestus* the best performing calibration area across all models and performance measures was the 100km radius around presence points, therefore this calibration area was selected as the most suitable for the mosquito vectors. By selecting absences within a 100km calibrating area it likely prevented absences being placed in areas without presences due to a lack of sampling effort or data availability, and ensured wide coverage for areas with limited presence and absence data but where sampling does occur across the country, such as the nationwide mosquito surveys in the UK.

*Table S. 2: A comparison of 5km, 50km and 100km calibration areas for Cx. pipiens and Cx. modestus. The performance metrics used for the comparison are True Positive Rate (TPR), True Negative Rate (TNR), Sorensen Index, Jaccard Index, F-measure on presence-background (FPB), omission rate (OR), True Skills Statistic (TSS), continuous Boyce index and inverse mean absolute error (IMAE). For all performance metrics a higher score indicates better model performance, and the majority are on a scale of 0 to 1, with the exceptions being FPB which uses 0 to 2 and TSS and Boyce which use -1 to 1.*

| ***Culex pipiens*** | | | | | | | | | | | | | | | | | | | | | | |
|---|---|---|---|---|---|---|---|---|---|---|---|---|---|---|---|---|---|---|---|---|---|---|
| **Calibration Area (m)** | **model** | **Thr value** | **TPR mean** | **TPR sd** | **TNR mean** | **TNR sd** | **SORENSEN mean** | **SORENSEN sd** | **JACCARD mean** | **JACCARD sd** | **FPB mean** | **FPB sd** | **OR mean** | **OR sd** | **TSS mean** | **TSS sd** | **AUC mean** | **AUC sd** | **BOYCE mean** | **BOYCE sd** | **IMAE mean** | **IMAE sd** |
| 5000 | SVM | 0.62 | 0.41 | 0.32 | 0.66 | 0.30 | 0.47 | 0.26 | 0.33 | 0.22 | 0.66 | 0.45 | 0.59 | 0.32 | 0.07 | 0.03 | 0.53 | 0.03 | 0.46 | 0.38 | 0.53 | 0.00 |
| | Random Forest | 0.64 | 0.07 | 0.03 | 0.96 | 0.00 | 0.13 | 0.06 | 0.07 | 0.03 | 0.14 | 0.07 | 0.93 | 0.03 | 0.03 | 0.04 | 0.48 | 0.00 | -0.15 | 0.27 | 0.52 | 0.00 |
| | NeuralNetwork | 0.60 | 0.47 | 0.26 | 0.63 | 0.21 | 0.53 | 0.19 | 0.37 | 0.18 | 0.75 | 0.36 | 0.53 | 0.26 | 0.09 | 0.05 | 0.53 | 0.04 | -0.16 | 0.07 | 0.53 | 0.00 |
| | MaxEnt | 0.63 | 0.48 | 0.26 | 0.61 | 0.24 | 0.54 | 0.17 | 0.38 | 0.16 | 0.76 | 0.32 | 0.52 | 0.26 | 0.09 | 0.02 | 0.53 | 0.00 | 0.42 | 0.18 | 0.53 | 0.00 |
| | GBM | 0.62 | 0.49 | 0.22 | 0.60 | 0.19 | 0.55 | 0.16 | 0.39 | 0.15 | 0.78 | 0.30 | 0.51 | 0.22 | 0.09 | 0.03 | 0.54 | 0.02 | 0.51 | 0.21 | 0.53 | 0.00 |
| | Ensemble mean | 0.58 | 0.44 | 0.47 | 0.62 | 0.45 | 0.45 | 0.38 | 0.33 | 0.32 | 0.65 | 0.64 | 0.56 | 0.47 | 0.06 | 0.02 | 0.52 | 0.02 | 0.56 | 0.27 | 0.53 | 0.00 |
| | Ensemble meanw | 0.59 | 0.47 | 0.26 | 0.63 | 0.22 | 0.53 | 0.19 | 0.37 | 0.18 | 0.75 | 0.36 | 0.53 | 0.26 | 0.09 | 0.04 | 0.54 | 0.04 | 0.43 | 0.29 | 0.53 | 0.00 |
| | Ensemble meansup | 0.60 | 0.46 | 0.24 | 0.64 | 0.22 | 0.53 | 0.17 | 0.37 | 0.16 | 0.73 | 0.32 | 0.54 | 0.24 | 0.09 | 0.02 | 0.54 | 0.03 | 0.29 | 0.16 | 0.53 | 0.00 |
| | Ensemble median | 0.60 | 0.72 | 0.17 | 0.34 | 0.17 | 0.67 | 0.06 | 0.51 | 0.07 | 1.02 | 0.14 | 0.28 | 0.17 | 0.06 | 0.00 | 0.52 | 0.01 | 0.52 | 0.47 | 0.53 | 0.00 |
| **Mean for performance evaluation across all models** | | | 0.45 | 0.11 | 0.63 | 0.11 | 0.49 | 0.07 | 0.35 | 0.06 | 0.69 | 0.12 | 0.55 | 0.11 | 0.08 | 0.02 | 0.53 | 0.01 | 0.32 | 0.10 | 0.53 | 0.00 |
| 50,000 | SVM | 0.34 | 0.58 | 0.14 | 0.76 | 0.06 | 0.57 | 0.12 | 0.40 | 0.12 | 0.81 | 0.24 | 0.42 | 0.14 | 0.34 | 0.08 | 0.71 | 0.07 | 0.83 | 0.04 | 0.58 | 0.02 |
| | Random Forest | 0.61 | 0.53 | 0.15 | 0.71 | 0.10 | 0.51 | 0.12 | 0.35 | 0.11 | 0.70 | 0.22 | 0.47 | 0.15 | 0.24 | 0.05 | 0.66 | 0.04 | 0.97 | 0.01 | 0.59 | 0.02 |
| | NeuralNetwork | 0.33 | 0.56 | 0.08 | 0.75 | 0.03 | 0.54 | 0.13 | 0.38 | 0.12 | 0.76 | 0.24 | 0.44 | 0.08 | 0.31 | 0.10 | 0.69 | 0.08 | 0.84 | 0.08 | 0.60 | 0.03 |
| | MaxEnt | 0.63 | 0.65 | 0.03 | 0.65 | 0.03 | 0.56 | 0.08 | 0.39 | 0.08 | 0.77 | 0.16 | 0.35 | 0.03 | 0.30 | 0.00 | 0.68 | 0.03 | 0.94 | 0.08 | 0.55 | 0.00 |
| | GBM | 0.36 | 0.60 | 0.26 | 0.69 | 0.18 | 0.55 | 0.15 | 0.38 | 0.15 | 0.77 | 0.30 | 0.40 | 0.26 | 0.29 | 0.08 | 0.68 | 0.07 | 0.85 | 0.01 | 0.58 | 0.00 |
| | Ensemble mean | 0.33 | 0.66 | 0.08 | 0.67 | 0.01 | 0.57 | 0.12 | 0.40 | 0.12 | 0.81 | 0.23 | 0.34 | 0.08 | 0.32 | 0.07 | 0.71 | 0.06 | 0.96 | 0.01 | 0.58 | 0.01 |

| | | | | | | | | | | | | | | | | | | | | | | |
|---|---|---|---|---|---|---|---|---|---|---|---|---|---|---|---|---|---|---|---|---|---|---|
| | Ensemble meanw | 0.33 | 0.64 | 0.06 | 0.69 | 0.01 | 0.57 | 0.11 | 0.40 | 0.11 | 0.80 | 0.22 | 0.36 | 0.06 | 0.33 | 0.06 | 0.71 | 0.06 | 0.95 | 0.01 | 0.58 | 0.01 |
| | Ensemble meansup | 0.31 | 0.55 | 0.09 | 0.78 | 0.01 | 0.55 | 0.12 | 0.39 | 0.11 | 0.77 | 0.23 | 0.45 | 0.09 | 0.33 | 0.08 | 0.71 | 0.07 | 0.95 | 0.01 | 0.59 | 0.02 |
| | Ensemble median | 0.26 | 0.58 | 0.16 | 0.76 | 0.07 | 0.56 | 0.13 | 0.40 | 0.13 | 0.80 | 0.26 | 0.42 | 0.16 | 0.33 | 0.09 | 0.70 | 0.08 | 0.89 | 0.11 | 0.58 | 0.01 |
| | **Mean for performance evaluation across all models** | | 0.59 | 0.04 | 0.72 | 0.05 | 0.55 | 0.01 | 0.39 | 0.01 | 0.78 | 0.02 | 0.41 | 0.04 | 0.31 | 0.02 | 0.69 | 0.01 | 0.91 | 0.05 | 0.58 | 0.01 |
| 100,000 | SVM | 0.21 | 0.70 | 0.10 | 0.71 | 0.05 | 0.50 | 0.21 | 0.35 | 0.19 | 0.70 | 0.38 | 0.30 | 0.10 | 0.42 | 0.14 | 0.75 | 0.10 | 0.70 | 0.18 | 0.69 | 0.01 |
| | Random Forest | 0.57 | 0.73 | 0.05 | 0.67 | 0.16 | 0.49 | 0.21 | 0.34 | 0.18 | 0.67 | 0.37 | 0.27 | 0.05 | 0.40 | 0.12 | 0.74 | 0.06 | 0.98 | 0.03 | 0.69 | 0.01 |
| | NeuralNetwork | 0.23 | 0.63 | 0.19 | 0.76 | 0.09 | 0.51 | 0.16 | 0.35 | 0.15 | 0.69 | 0.30 | 0.37 | 0.19 | 0.39 | 0.10 | 0.73 | 0.08 | 0.87 | 0.04 | 0.71 | 0.01 |
| | MaxEnt | 0.63 | 0.72 | 0.08 | 0.67 | 0.03 | 0.49 | 0.16 | 0.33 | 0.14 | 0.67 | 0.28 | 0.28 | 0.08 | 0.40 | 0.05 | 0.74 | 0.06 | 0.96 | 0.06 | 0.60 | 0.00 |
| | GBM | 0.26 | 0.60 | 0.20 | 0.80 | 0.06 | 0.51 | 0.19 | 0.35 | 0.17 | 0.71 | 0.34 | 0.40 | 0.20 | 0.40 | 0.14 | 0.75 | 0.09 | 0.79 | 0.24 | 0.71 | 0.03 |
| | Ensemble mean | 0.25 | 0.65 | 0.20 | 0.77 | 0.10 | 0.52 | 0.16 | 0.36 | 0.15 | 0.72 | 0.30 | 0.35 | 0.20 | 0.42 | 0.11 | 0.76 | 0.08 | 0.95 | 0.02 | 0.68 | 0.01 |
| | Ensemble meanw | 0.25 | 0.65 | 0.21 | 0.77 | 0.10 | 0.52 | 0.17 | 0.36 | 0.15 | 0.72 | 0.30 | 0.35 | 0.21 | 0.42 | 0.11 | 0.76 | 0.08 | 0.95 | 0.02 | 0.68 | 0.01 |
| | Ensemble meansup | 0.12 | 0.70 | 0.10 | 0.71 | 0.05 | 0.50 | 0.21 | 0.35 | 0.19 | 0.70 | 0.38 | 0.30 | 0.10 | 0.42 | 0.14 | 0.75 | 0.10 | 0.70 | 0.18 | 0.69 | 0.01 |
| | Ensemble median | 0.16 | 0.64 | 0.19 | 0.78 | 0.06 | 0.52 | 0.19 | 0.36 | 0.17 | 0.73 | 0.34 | 0.36 | 0.19 | 0.42 | 0.13 | 0.76 | 0.08 | 0.89 | 0.11 | 0.69 | 0.01 |
| | **Mean for performance evaluation across all models** | | 0.67 | 0.04 | 0.74 | 0.05 | 0.51 | 0.01 | 0.35 | 0.01 | 0.70 | 0.02 | 0.33 | 0.04 | 0.41 | 0.01 | 0.75 | 0.01 | 0.86 | 0.11 | 0.68 | 0.04 |
| **Highest scoring Calibration Area** | | | **CA100000** | | **CA100000** | | **CA50000** | | **CA50000** | | **CA50000** | | **CA5000** | | **CA100000** | | **CA100000** | | **CA50000** | | **CA100000** | |
| **Selected model based on individual performance metric for CA100000** | | | **Random Forest** | | **Ensemble median** | | **Ensemble meanw** | | **Ensemble meanw** | | **Ensemble median** | | **GBM** | | **Ensemble meanw** | | **Ensemble meanw** | | **Random forest** | | **Neural network** | |
| ***Culex modestus*** | | | | | | | | | | | | | | | | | | | | | | |

| Calibration area | model | thr value | TPR mean | TPR sd | TNR mean | TNR sd | SORENSEN mean | SORENSEN sd | JACCARD mean | JACCARD sd | FPB mean | FPB sd | OR mean | OR sd | TSS mean | TSS sd | AUC mean | AUC sd | BOYCE mean | BOYCE sd | IMAE mean | IMAE sd |
|---|---|---|---|---|---|---|---|---|---|---|---|---|---|---|---|---|---|---|---|---|---|---|
| 5000 | SVM | 0.67 | 0.34 | 0.04 | 0.81 | 0.08 | 0.43 | 0.01 | 0.27 | 0.01 | 0.54 | 0.02 | 0.66 | 0.04 | 0.15 | 0.03 | 0.51 | 0.05 | 0.74 | 0.04 | 0.52 | 0.01 |
| | Random Forest | 0.64 | 0.81 | 0.18 | 0.37 | 0.31 | 0.61 | 0.01 | 0.43 | 0.01 | 0.87 | 0.02 | 0.19 | 0.18 | 0.19 | 0.13 | 0.52 | 0.07 | 0.59 | 0.42 | 0.51 | 0.02 |
| | NeuralNetwork | 0.45 | 0.77 | 0.24 | 0.39 | 0.30 | 0.58 | 0.04 | 0.41 | 0.04 | 0.82 | 0.09 | 0.23 | 0.24 | 0.16 | 0.05 | 0.53 | 0.05 | 0.84 | 0.18 | 0.51 | 0.00 |
| | MaxEnt | 0.63 | 0.59 | 0.04 | 0.63 | 0.01 | 0.56 | 0.04 | 0.39 | 0.03 | 0.78 | 0.07 | 0.41 | 0.04 | 0.22 | 0.05 | 0.57 | 0.04 | 0.72 | 0.03 | 0.53 | 0.05 |
| | GBM | 0.46 | 0.33 | 0.07 | 0.79 | 0.08 | 0.40 | 0.04 | 0.25 | 0.03 | 0.50 | 0.06 | 0.67 | 0.07 | 0.11 | 0.02 | 0.50 | 0.02 | 0.35 | 0.43 | 0.51 | 0.01 |
| | Ensemble mean | 0.50 | 0.33 | 0.02 | 0.79 | 0.04 | 0.40 | 0.01 | 0.25 | 0.00 | 0.51 | 0.01 | 0.67 | 0.02 | 0.11 | 0.02 | 0.53 | 0.01 | 0.30 | 0.03 | 0.52 | 0.02 |
| | Ensemble meanw | 0.52 | 0.61 | 0.11 | 0.55 | 0.07 | 0.54 | 0.05 | 0.37 | 0.05 | 0.75 | 0.10 | 0.39 | 0.11 | 0.16 | 0.04 | 0.54 | 0.02 | 0.39 | 0.02 | 0.52 | 0.02 |
| | Ensemble meansup | 0.58 | 0.48 | 0.07 | 0.69 | 0.12 | 0.50 | 0.01 | 0.34 | 0.00 | 0.67 | 0.01 | 0.52 | 0.07 | 0.17 | 0.06 | 0.56 | 0.05 | 0.53 | 0.20 | 0.52 | 0.03 |
| | Ensemble median | 0.40 | 0.63 | 0.27 | 0.50 | 0.33 | 0.53 | 0.06 | 0.36 | 0.06 | 0.72 | 0.12 | 0.38 | 0.27 | 0.12 | 0.06 | 0.54 | 0.04 | 0.64 | 0.29 | 0.51 | 0.01 |
| | **Mean performance across all models** | | 0.54 | 0.14 | 0.61 | 0.12 | 0.51 | 0.07 | 0.34 | 0.06 | 0.68 | 0.12 | 0.46 | 0.14 | 0.16 | 0.04 | 0.53 | 0.03 | 0.57 | 0.17 | 0.52 | 0.01 |
| 50,000 | SVM | 0.25 | 0.67 | 0.13 | 0.65 | 0.11 | 0.50 | 0.00 | 0.33 | 0.00 | 0.66 | 0.01 | 0.33 | 0.13 | 0.32 | 0.02 | 0.64 | 0.03 | 0.96 | 0.05 | 0.62 | 0.01 |
| | Random Forest | 0.59 | 0.76 | 0.04 | 0.51 | 0.11 | 0.48 | 0.05 | 0.32 | 0.04 | 0.64 | 0.09 | 0.24 | 0.04 | 0.27 | 0.07 | 0.59 | 0.05 | 0.96 | 0.00 | 0.64 | 0.02 |
| | NeuralNetwork | 0.24 | 0.88 | 0.17 | 0.49 | 0.03 | 0.52 | 0.11 | 0.36 | 0.10 | 0.72 | 0.21 | 0.12 | 0.17 | 0.36 | 0.21 | 0.66 | 0.07 | 0.94 | 0.05 | 0.61 | 0.01 |
| | MaxEnt | 0.63 | 0.71 | 0.41 | 0.44 | 0.62 | 0.44 | 0.06 | 0.28 | 0.05 | 0.57 | 0.10 | 0.29 | 0.41 | 0.15 | 0.21 | 0.45 | 0.13 | 0.84 | 0.15 | 0.47 | 0.04 |
| | GBM | 0.33 | 0.51 | 0.24 | 0.80 | 0.08 | 0.48 | 0.14 | 0.32 | 0.12 | 0.64 | 0.24 | 0.49 | 0.24 | 0.31 | 0.16 | 0.59 | 0.11 | 0.94 | 0.02 | 0.66 | 0.04 |
| | Ensemble mean | 0.33 | 0.37 | 0.23 | 0.83 | 0.09 | 0.38 | 0.16 | 0.24 | 0.12 | 0.48 | 0.24 | 0.63 | 0.23 | 0.19 | 0.14 | 0.53 | 0.07 | 0.88 | 0.12 | 0.60 | 0.00 |
| | Ensemble meanw | 0.29 | 0.63 | 0.14 | 0.58 | 0.28 | 0.45 | 0.07 | 0.29 | 0.06 | 0.58 | 0.12 | 0.37 | 0.14 | 0.21 | 0.14 | 0.56 | 0.06 | 0.93 | 0.06 | 0.62 | 0.00 |
| | Ensemble meansup | 0.22 | 0.64 | 0.36 | 0.66 | 0.23 | 0.47 | 0.11 | 0.31 | 0.09 | 0.62 | 0.19 | 0.36 | 0.36 | 0.29 | 0.13 | 0.62 | 0.07 | 0.95 | 0.06 | 0.63 | 0.00 |

| | | | | | | | | | | | | | | | | | | | | | | |
|---|---|---|---|---|---|---|---|---|---|---|---|---|---|---|---|---|---|---|---|---|---|---|
| | Ensemble median | 0.27 | 0.800 | 0.059 | 0.502 | 0.011 | 0.496 | 0.053 | 0.330 | 0.047 | 0.661 | 0.094 | 0.200 | 0.059 | 0.302 | 0.070 | 0.633 | 0.068 | 0.988 | 0.003 | 0.616 | 0.009 |
| | **Mean performance across all models** | | 0.66 | 0.15 | 0.61 | 0.16 | 0.47 | 0.04 | 0.31 | 0.03 | 0.62 | 0.07 | 0.34 | 0.15 | 0.27 | 0.07 | 0.59 | 0.07 | 0.93 | 0.05 | 0.61 | 0.07 |
| 100,000 | SVM | 0.22 | 0.68 | 0.12 | 0.66 | 0.11 | 0.50 | 0.02 | 0.33 | 0.02 | 0.67 | 0.04 | 0.32 | 0.12 | 0.34 | 0.01 | 0.66 | 0.02 | 0.89 | 0.07 | 0.64 | 0.02 |
| | Random Forest | 0.62 | 0.32 | 0.18 | 0.93 | 0.04 | 0.40 | 0.17 | 0.26 | 0.13 | 0.52 | 0.26 | 0.68 | 0.18 | 0.25 | 0.14 | 0.56 | 0.08 | 0.72 | 0.15 | 0.67 | 0.00 |
| | NeuralNetwork | 0.33 | 0.79 | 0.17 | 0.61 | 0.16 | 0.53 | 0.03 | 0.36 | 0.03 | 0.72 | 0.06 | 0.21 | 0.17 | 0.39 | 0.01 | 0.68 | 0.08 | 0.93 | 0.08 | 0.67 | 0.02 |
| | MaxEnt | 0.63 | 0.63 | 0.21 | 0.70 | 0.09 | 0.50 | 0.10 | 0.33 | 0.09 | 0.67 | 0.18 | 0.37 | 0.21 | 0.33 | 0.12 | 0.61 | 0.08 | 0.73 | 0.05 | 0.54 | 0.10 |
| | GBM | 0.28 | 0.58 | 0.19 | 0.76 | 0.16 | 0.50 | 0.00 | 0.33 | 0.00 | 0.67 | 0.00 | 0.42 | 0.19 | 0.34 | 0.03 | 0.63 | 0.03 | 0.95 | 0.06 | 0.71 | 0.04 |
| | Ensemble mean | 0.29 | 0.51 | 0.18 | 0.83 | 0.08 | 0.50 | 0.09 | 0.34 | 0.08 | 0.67 | 0.17 | 0.49 | 0.18 | 0.34 | 0.10 | 0.67 | 0.06 | 0.92 | 0.07 | 0.64 | 0.03 |
| | Ensemble meanw | 0.27 | 0.58 | 0.28 | 0.74 | 0.20 | 0.49 | 0.08 | 0.33 | 0.07 | 0.65 | 0.14 | 0.42 | 0.28 | 0.32 | 0.08 | 0.67 | 0.05 | 0.91 | 0.10 | 0.64 | 0.04 |
| | Ensemble meansup | 0.27 | 0.57 | 0.13 | 0.75 | 0.09 | 0.49 | 0.05 | 0.33 | 0.05 | 0.65 | 0.09 | 0.43 | 0.13 | 0.32 | 0.04 | 0.67 | 0.05 | 0.92 | 0.10 | 0.64 | 0.04 |
| | Ensemble median | 0.24 | 0.64 | 0.09 | 0.72 | 0.06 | 0.52 | 0.04 | 0.35 | 0.04 | 0.70 | 0.08 | 0.36 | 0.09 | 0.36 | 0.03 | 0.67 | 0.01 | 0.95 | 0.04 | 0.66 | 0.02 |
| | **Mean performance across all models** | | 0.59 | 0.05 | 0.74 | 0.04 | 0.49 | 0.01 | 0.33 | 0.01 | 0.66 | 0.02 | 0.41 | 0.05 | 0.33 | 0.02 | 0.65 | 0.03 | 0.88 | 0.08 | 0.65 | 0.05 |
| **Highest scoring Calibration Area** | | | **CA50000** | | **CA100000** | | **CA5000** | | **CA5000** | | **CA5000** | | **CA5000** | | **CA10000** | | **CA100000** | | **CA50000** | | **CA100000** | |
| **Selected model based on individual performance metric for CA100000** | | | **Nueral Network** | | **Random forest** | | **Neural Network** | | **Neural Network** | | **Neural Networ** | | **Random Forest** | | **Neural Network** | | **Neural Network** | | **GBM / Ensemble median** | | **GBM** | |

### SI. 3: Sensitivity analysis of CLIMEX modelling

*Table S. 3: Sensitivity analysis was carried out in CLIMEX for the Cx. pipiens model and the results are shown below for the impact of the parameters initially considered on the Ecoclimatic Index output.*

| Parameter | Model parameter | Low | High | Ecoclimatic Index (EI) Change |
|---|---|---|---|---|
| Limiting low moisture (SM0) | 0.25 | 0.15 | 0.35 | 4.45 |
| Lower optimal moisture (SM1) | 0.8 | 0.7 | 0.9 | 2.87 |
| Upper optimal moisture (SM2) | 1.5 | 1.4 | 1.6 | 0.06 |
| Limiting high moisture (SM3) | 2.5 | 2.4 | 2.6 | 0.01 |
| Limiting low temperature (DV0) | 7 | 6 | 8 | 0.81 |
| Lower optimal temperature (DV1) | 15 | 14 | 16 | 1.75 |
| Upper optimal temperature (DV2) | 25 | 24 | 26 | 0.39 |
| Limiting high temperature (DV3) | 29 | 28 | 30 | 0.20 |
| Diapause Induction Daylength (DPD0) | 13.5 | 13 | 14 | 1.27 |
| Diapause Induction Temperature (DPT0) | 15 | 14 | 16 | 0.45 |
| Diapause Termination Temperature (DPT1) | 6.5 | 5.5 | 7.5 | 3.32 |
| Diapause Development Days (DPD) | -100 | -120 | -80 | 0.00 |
| Degree-days per Generation (PDD) | 83 | 66.4 | 99.6 | 0 |

### SI. 4: Partial dependence plots for vector models

For *Cx. pipiens*, the partial-dependence plots showed that for many variables the test data were within the ranges of the training data, though some variables required extrapolation into the novel regions such as the mean diurnal range (bio2) and precipitation during the driest month (bio14) (Figure S. 2). Compared to *Cx. pipiens,* much more of the UK had to be projected for *Cx. modestus,* likely due to the sample size being considerably smaller (Figure S. 3). Overall, when projecting the models, they generally followed the trend of the last known values of the full range, however there were high levels of variation between the model approaches. For example, for precipitation during the driest month (bio14) in the RF, MaxEnt and GBM models the suitability remained relatively constant into the projected range however for the Net and SVM models there was greater suitability predicted once outside of the range of the training dataset.

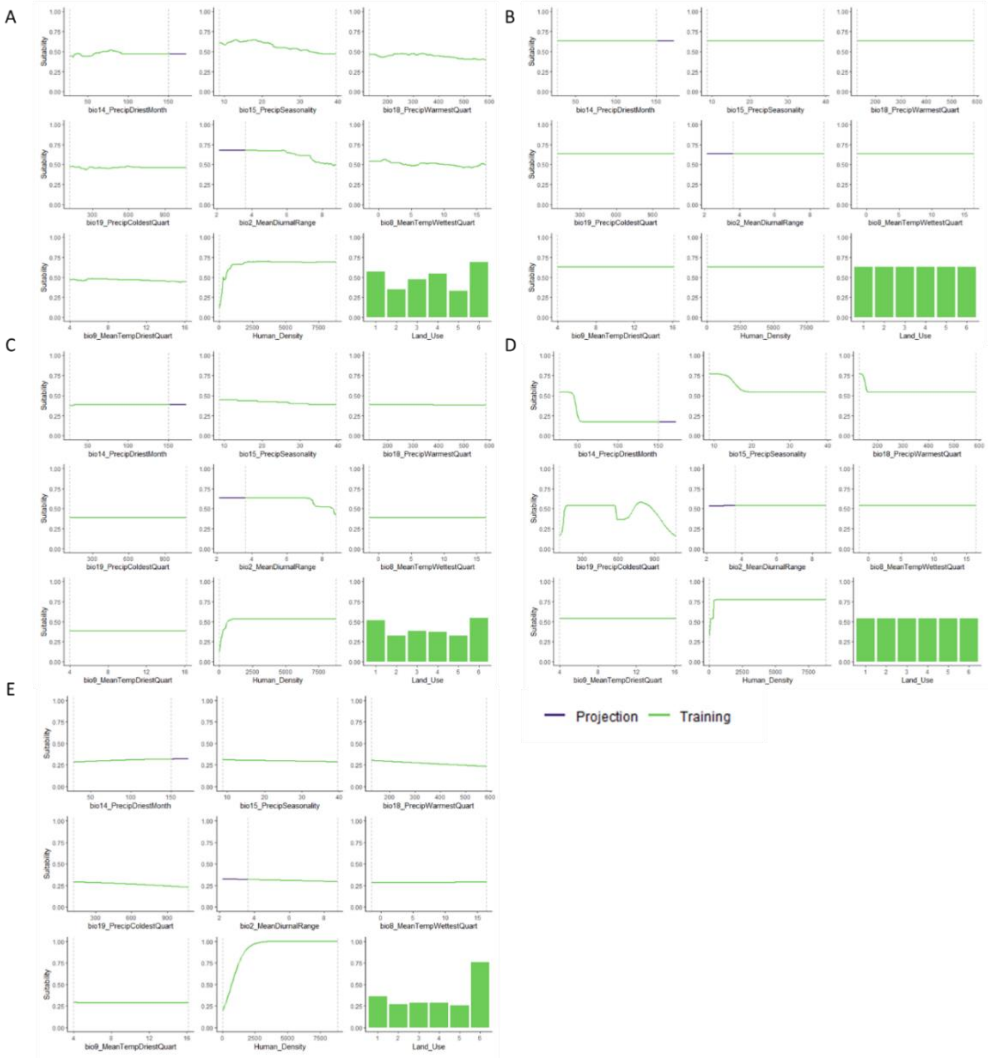


*Figure S. 2: Partial dependence plots of the training and projection for the five independent correlative models for Cx. pipiens, A) Random Forest, B) MaxEnt, C) Generalized Boosted Regression, D) Neural Network and E) Support Vector Machine. Values within the range of the training dataset are shown in green whereas values outside of the training dataset but within the projection area are shown in purple.*

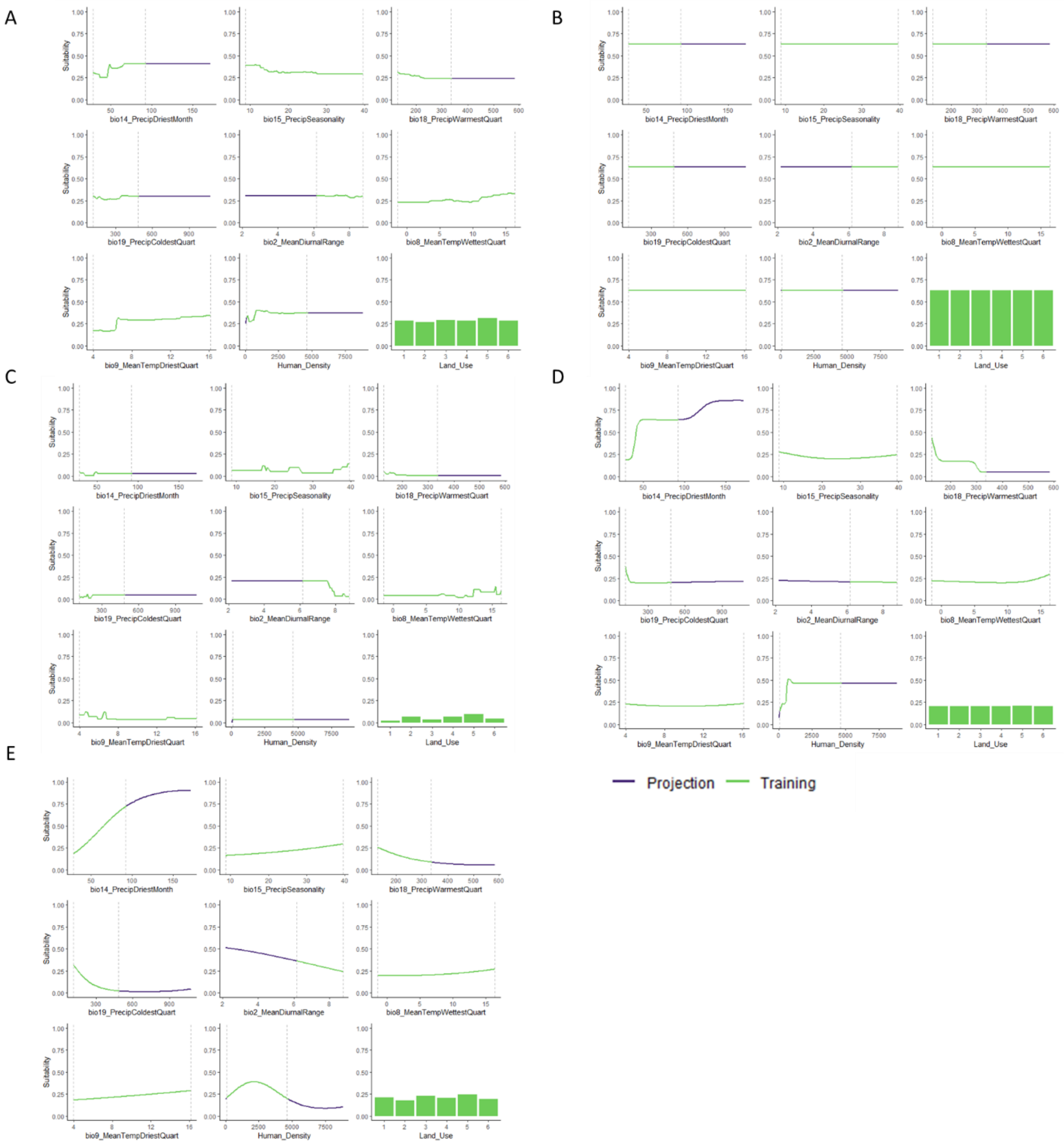


*Figure S. 3: Partial dependence plots of the training and projection for the five independent correlative models for Cx. modestus, A) Random Forest, B) MaxEnt, C) Generalized Boosted Regression, D) Neural Network and E) Support Vector Machine. Values within the range of the training dataset are shown in green whereas values outside of the training dataset but within the projection area are shown in purple.*

### SI. 5: WNV host species distribution modelling

Avian host species distribution models were produced, representing corvid and non-corvid Passeriformes. Both models performed well overall and the highest scoring model was selected by taking into account the model that most frequently performed highest across the performance metrics. For corvids, this was the GBM model, whilst the mean-weighted ensemble model performed best for non-corvids (Table S. 4). The distribution of both avian host groups was very similar, with much of the UK considered to be highly suitable, such as the south-east, south-west, west midlands and north-west (Figure S. 2). This pattern of suitability remained consistent in future predictions for both climate change scenarios (SSP1 and SSP5), with general suitability increasing across the UK in 2100 for both corvid and non-corvid Passeriformes, though some areas do also show a reduction in suitability when predicting into the future (Figure S. 2).

*Table S. 4: Model performance scores of the avian host models. The performance metrics are True positive rate (TPR), True negative rate (TNR), Sorensen, Jaccard, F-measure (FPB), Omission rate (OR), True skill statistic (TSS), Area under curve (AUC), Boyce index and Inverse mean absolute error (IMAE). Higher values indicate better model performance.*

| | TPR | | TNR | | Sorensen | | Jaccard | | FPB | | OR | | TSS | | SUC | | Boyce | | IMAE | |
|---|---|---|---|---|---|---|---|---|---|---|---|---|---|---|---|---|---|---|---|---|
| **score range** | **0 to 1** | | **0 to 1** | | **0 to 1** | | **0 to 1** | | **0 to 2** | | **0 to 1** | | **-1 to 1** | | **0 to 1** | | **-1 to 1** | | **0 to 1** | |
| | mean | sd | mean | sd | mean | sd | mean | sd | mean | Sd | mean | sd | mean | sd | mean | sd | mean | sd | mean | sd |
| *Corvid* | | | | | | | | | | | | | | | | | | | | |
| **Neural Network** | 0.78 | 0.05 | 0.81 | 0.00 | 0.79 | 0.01 | 0.65 | 0.01 | 1.31 | 0.03 | 0.22 | 0.05 | 0.59 | 0.05 | 0.85 | 0.03 | 0.98 | 0.02 | 0.70 | 0.02 |
| **GBM** | 0.77 | 0.05 | 0.82 | 0.04 | 0.79 | 0.02 | 0.65 | 0.03 | 1.31 | 0.07 | 0.23 | 0.05 | 0.59 | 0.08 | 0.86 | 0.04 | 0.99 | 0.01 | 0.70 | 0.02 |
| **Random Forest** | 0.73 | 0.05 | 0.83 | 0.01 | 0.77 | 0.02 | 0.62 | 0.03 | 1.24 | 0.06 | 0.27 | 0.05 | 0.56 | 0.07 | 0.86 | 0.03 | 0.99 | 0.01 | 0.68 | 0.00 |
| **Ens mean weighted** | 0.75 | 0.06 | 0.87 | 0.02 | 0.80 | 0.03 | 0.66 | 0.04 | 1.33 | 0.08 | 0.25 | 0.06 | 0.62 | 0.08 | 0.87 | 0.04 | 1.00 | 0.01 | 0.69 | 0.01 |
| **Ens mean** | 0.75 | 0.06 | 0.87 | 0.02 | 0.80 | 0.03 | 0.66 | 0.04 | 1.33 | 0.08 | 0.25 | 0.06 | 0.62 | 0.08 | 0.87 | 0.04 | 1.00 | 0.01 | 0.69 | 0.01 |
| **Ens meanw** | 0.78 | 0.07 | 0.82 | 0.01 | 0.79 | 0.02 | 0.66 | 0.02 | 1.31 | 0.05 | 0.22 | 0.07 | 0.60 | 0.06 | 0.86 | 0.03 | 0.99 | 0.02 | 0.70 | 0.02 |
| **Ens meansup** | 0.77 | 0.08 | 0.85 | 0.00 | 0.80 | 0.03 | 0.66 | 0.04 | 1.32 | 0.08 | 0.23 | 0.08 | 0.61 | 0.08 | 0.87 | 0.03 | 0.99 | 0.01 | 0.70 | 0.02 |
| *Non-corvid* | | | | | | | | | | | | | | | | | | | | |
| **Neural Network** | 0.82 | 0.02 | 0.80 | 0.04 | 0.81 | 0.05 | 0.68 | 0.06 | 1.36 | 0.13 | 0.18 | 0.02 | 0.62 | 0.06 | 0.88 | 0.03 | 1.00 | 0.00 | 0.72 | 0.01 |
| **GBM** | 0.81 | 0.06 | 0.82 | 0.01 | 0.81 | 0.06 | 0.68 | 0.08 | 1.37 | 0.17 | 0.19 | 0.06 | 0.63 | 0.07 | 0.88 | 0.03 | 1.00 | 0.00 | 0.73 | 0.02 |
| **Random Forest** | 0.85 | 0.03 | 0.84 | 0.02 | 0.85 | 0.04 | 0.74 | 0.06 | 1.47 | 0.11 | 0.15 | 0.03 | 0.70 | 0.04 | 0.92 | 0.01 | 1.00 | 0.00 | 0.77 | 0.01 |
| **Ens mean weighted** | 0.84 | 0.03 | 0.84 | 0.03 | 0.84 | 0.05 | 0.72 | 0.07 | 1.44 | 0.13 | 0.16 | 0.03 | 0.68 | 0.06 | 0.91 | 0.02 | 1.00 | 0.00 | 0.74 | 0.01 |
| **Ens mean** | 0.84 | 0.03 | 0.84 | 0.02 | 0.84 | 0.05 | 0.72 | 0.07 | 1.45 | 0.13 | 0.16 | 0.03 | 0.68 | 0.06 | 0.91 | 0.02 | 1.00 | 0.00 | 0.74 | 0.01 |
| **Ens meanw** | 0.85 | 0.03 | 0.84 | 0.02 | 0.85 | 0.04 | 0.74 | 0.06 | 1.47 | 0.11 | 0.15 | 0.03 | 0.70 | 0.04 | 0.92 | 0.01 | 1.00 | 0.00 | 0.77 | 0.01 |
| **Ens meansup** | 0.83 | 0.05 | 0.83 | 0.01 | 0.83 | 0.05 | 0.71 | 0.08 | 1.42 | 0.15 | 0.17 | 0.05 | 0.67 | 0.06 | 0.90 | 0.02 | 1.00 | 0.00 | 0.74 | 0.01 |

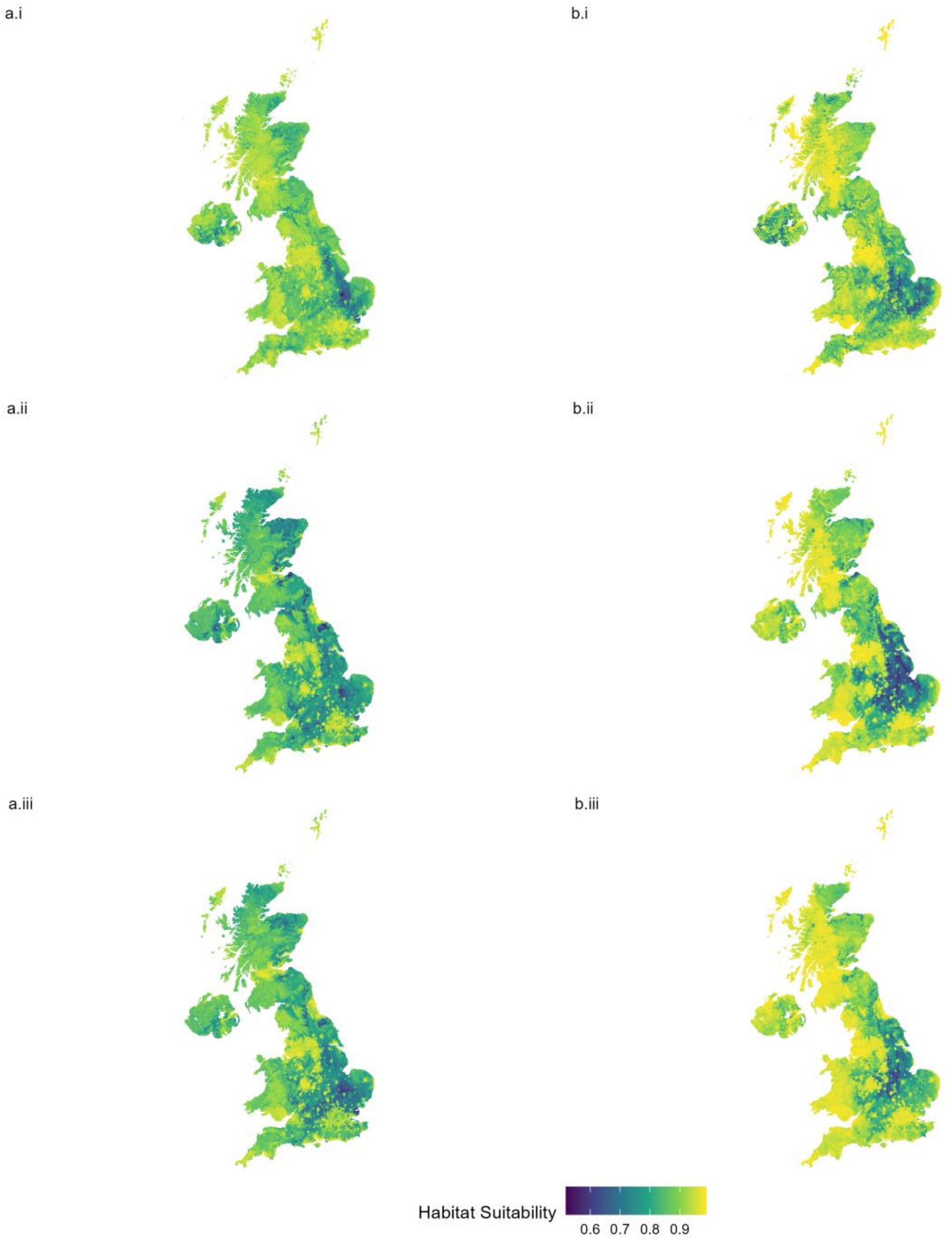


*Figure S. 4: Host species distribution models for a) corvid and b) non-corvid representative groups, for i) current, ii) SSP1 2100 and iii) SSP5 2100 predictions.*
